# Integrating analog and digital modes of gene expression at Arabidopsis *FLC*

**DOI:** 10.1101/2022.07.04.498694

**Authors:** Rea L. Antoniou-Kourounioti, Anis Meschichi, Svenja Reeck, Scott Berry, Govind Menon, Yusheng Zhao, John A. Fozard, Terri L. Holmes, Huamei Wang, Matthew Hartley, Caroline Dean, Stefanie Rosa, Martin Howard

## Abstract

Quantitative gene regulation at the cell population-level can be achieved by two fundamentally different modes of regulation at individual gene copies. A “digital” mode involves binary ON/OFF expression states, with population-level variation arising from the proportion of gene copies in each state, while an “analog” mode involves graded expression levels at each gene copy. At the Arabidopsis floral repressor *FLOWERING LOCUS C (FLC),* “digital” Polycomb silencing is known to facilitate quantitative epigenetic memory in response to cold. However, whether *FLC* regulation before cold involves analog or digital modes is unknown. Using quantitative fluorescent imaging of *FLC* mRNA and protein, together with mathematical modelling, we find that *FLC* expression before cold is regulated by both analog and digital modes. We observe a temporal separation between the two modes, with analog preceding digital. The analog mode can maintain intermediate expression levels at individual *FLC* gene copies, before subsequent digital silencing, consistent with the copies switching OFF stochastically and heritably without cold. This switch leads to a slow reduction in *FLC* expression at the cell population-level. These data present a new paradigm for gradual repression, elucidating how analog transcriptional and digital epigenetic memory pathways can be integrated.

## Introduction

One of the most fundamental questions in molecular biology is how quantitative gene expression is achieved. Traditionally, such regulation is ascribed to sequence-specific transcription factors that bind to regulatory DNA elements. According to the concentration of the transcription factors, gene expression can then be quantitatively up or down-regulated. While such regulation undoubtedly occurs in many systems, it has become abundantly clear in recent years that this paradigm is fundamentally incomplete. This is especially so in eukaryotes where quantitative transcriptional regulation can arise from modulation of the local chromatin environment of a gene. For example, by varying the type and level of histone modifications, DNA accessibility can be radically altered (Ahmad, Henikoff et al. 2022). In one scenario, nucleosome positioning affects the ability of transcription factors to bind. Another possibility is that alteration of the chromatin environment directly affects the kinetics of transcription (Coulon, Ferguson et al. 2014) by altering how fast the RNA polymerase elongates.

For the Arabidopsis floral repressor gene *FLOWERING LOCUS C* (*FLC*), it has been shown that expression levels are quantitatively reduced by a prolonged duration of cold. This quantitative response is achieved through individual *FLC* gene copies making a cis-mediated, digital switch from an “ON” (expressing) state to an “OFF” (silenced) state. This switch is asynchronous between loci, even in the same cell, with the number of switched OFF gene copies increasing over time in the cold. This results in a gradual decrease of *FLC* expression over time at a whole plant level (Angel, Song et al. 2011) with silenced gene copies covered by high levels of the silencing histone mark H3K27me3 controlled by the Polycomb system through Polycomb Repressive Complex 2 (PRC2). This mode of regulation is called “digital”, to highlight the discrete ON/OFF states for each gene copy (Fig. 1A – Digital Regulation, (Munsky and Neuert 2015)). Such digital regulation has also been observed in many other systems, both natural (Saxton and Rine 2022) and engineered (Bintu, Yong et al. 2016).

**Figure 1.**
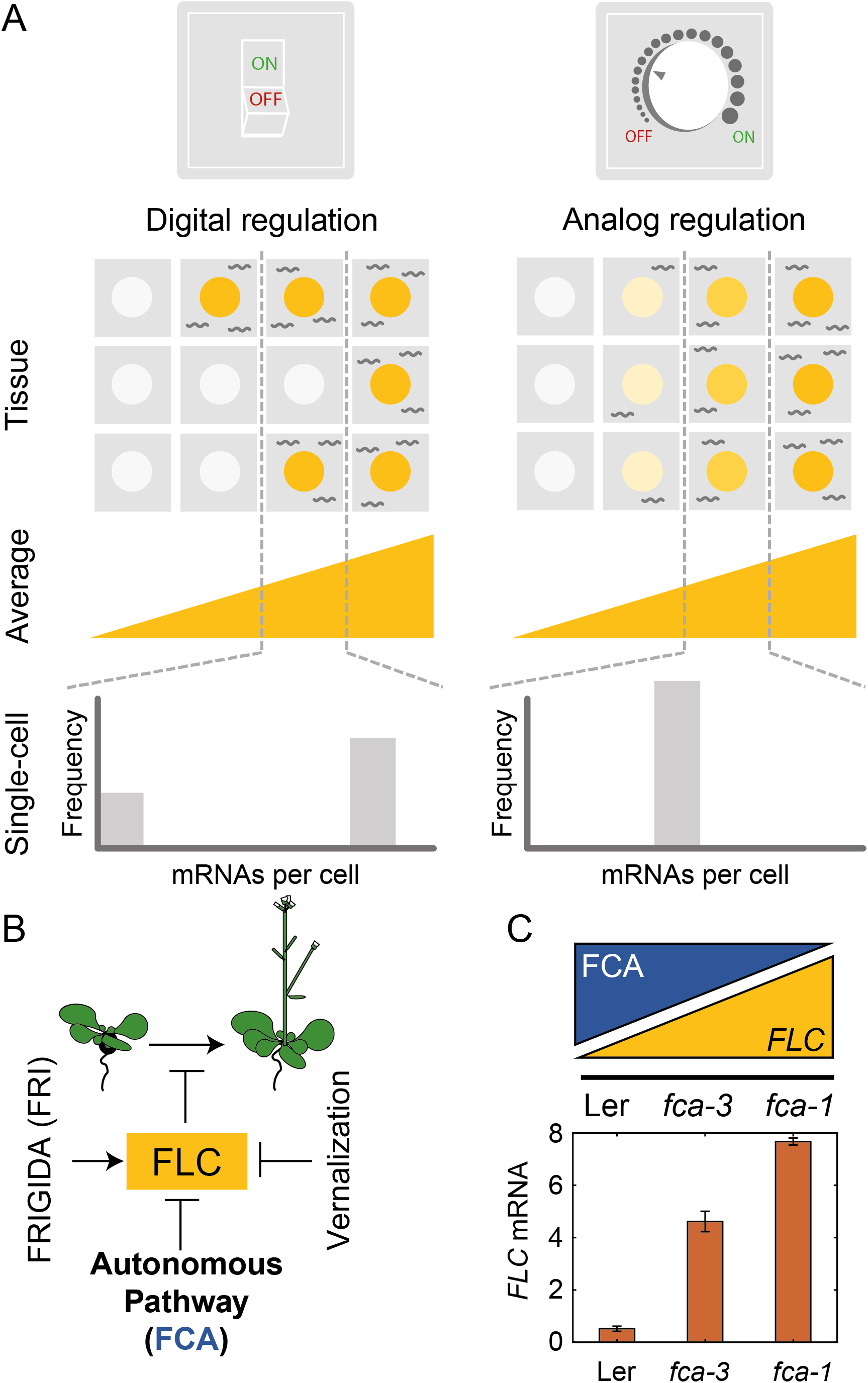
Schematic of digital and analog gene regulation. (A) Digital regulation (left) corresponds to loci being in an ‘ON’ state (yellow) or ‘OFF’ state (white), where we assume for simplicity that there is only one gene copy per cell. At the tissue level, moving from low to high average expression (columns left to right) is achieved by a change in the fraction of cells in each of the two states. This mode is distinct from analog regulation (right), where each cell has a graded expression level that roughly corresponds to the overall population average. (B) *FLC* represses the transition to flowering and is controlled by *FRIGIDA*, the autonomous pathway and the vernalization pathway (inducing digital epigenetic silencing in the cold). (C) Reduced *FCA* activity leads to higher cell population-level *FLC* expression in *fca* mutants. Wild-type Ler has lowest *FLC* expression, *fca-3* intermediate and *fca-1* highest. Expression is measured by qPCR relative to the house-keeping gene index (geometric mean of *PP2A* and *UBC)*. Error bars show SEM of n=3 biological replicates measured 7 days after sowing.

An alternative mode of quantitative gene regulation is one that allows graded expression levels to be maintained at each gene copy, rather than just an ON or an OFF state. Quantitative regulation at the cell population-level can then be achieved by tuning the expression level uniformly at all gene copies, as in inducible gene expression systems. A well-characterized example of such behaviour is in the level of stress-responsive gene expression as controlled by the transcription factor Msn2 in budding yeast (Stewart-Ornstein, Nelson et al. 2013). This graded mode of regulation is called “analog” (Fig. 1A – Analog Regulation, (Munsky and Neuert 2015)), in contrast to the digital alternative.

While *FLC* loci are known to have digital behaviour during and after a cold treatment, allowing them to robustly hold epigenetic memory of cold exposure, it remains unknown how the starting expression levels (prior to cold) are regulated quantitatively in terms of analog versus digital control. After plants germinate, they grow as seedlings that have not experienced any cold exposure, or digital Polycomb switching, so quantitative variation in *FLC* expression seen in young seedlings could represent different cellular proportions of digitally regulated *FLC*, as well as graded transcriptional changes. More generally, elucidating the interplay between analog and digital control is essential for a more in-depth understanding of quantitative gene regulation. Although digital and analog modes of repression have been separately studied in the past (see, e.g. (Munsky and Neuert 2015)), how these two fundamentally different modes of regulation might be combined has not been considered. *FLC* is an ideal system to study this question, due to its digital control after cold, as well as the wealth of knowledge about its regulation at all stages (Berry and Dean 2015, Wu, Fang et al. 2019).

*FLC* levels are set during embryogenesis by competition between an *FLC* activator called *FRIGIDA* (*FRI*) and the so-called autonomous repressive pathway (Fig. 1B, (Li, Jiang et al. 2018, Schon, Baxter et al. 2021)). The commonly used Arabidopsis accessions Ler and Col-0, have mutations in the *FRI* gene, allowing the autonomous pathway to dominate, thereby repressing *FLC* during vegetative development and resulting in a rapidly cycling summer annual lifestyle (Johanson, West et al. 2000). On the other hand, introgressing an active *FRI* allele (Col*FRI* (Lee and Amasino 1995)), generates an initially high *FLC* expression state, which then requires cold for *FLC* repression and subsequent flowering. However, both these cases, with high (Col*FRI*) or low (Ler, Col-0) *FLC* expression, are extreme examples, where essentially all *FLC* loci are either expressed to a high level or not, before cold. It is therefore difficult to use these genotypes to understand any possible interplay between digital and analog control, as only the extremes are exhibited. Instead, what is required is a genotype with intermediate *FLC* expression at a cell population-level during embryogenesis. Such a genotype at a single gene copy level might exhibit either a graded (analog) or all-or-nothing (digital) *FLC* expression before cold, thereby allowing dissection of whether analog or digital regulation is at work.

In this work, we exploited two mutant alleles in *FLOWERING CONTROL LOCUS A* (*FCA*), which is part of the *FLC-*repressive autonomous pathway, thereby systematically varying overall *FLC* levels. One of these mutants, *fca-1* (Koornneef, Hanhart et al. 1991), is a complete loss of function and therefore exhibits late flowering (Fig. 1S1A) and high *FLC* expression before cold (Fig. 1C, 1S1B-D), similar to Col*FRI*. The wildtype, Ler, has a fully functional autonomous pathway and so exhibits low *FLC* levels before cold (Fig. 1C, 1S1B-D) and early flowering (Fig. 1S1A), with the *FLC* gene covered by the silencing histone mark H3K27me3 (Wu, Fang et al. 2019). Crucially, however, the *fca-3* mutant (Koornneef, Hanhart et al. 1991) displays intermediate cell population-level *FLC* expression (Fig. 1C, 1S1B-D) and flowering time (Fig. 1S1A). This property allows us to systematically dissect the interplay of analog and digital regulation at *FLC* before cold using a combination of single cell and whole plant assays, together with mathematical modelling, revealing a temporal separation between the two regulatory modes.

## Results

### Analysis of *fca* alleles reveals both analog and digital regulation at *FLC*

To investigate the mode of repression arising from regulation by the autonomous pathway, we utilized the *fca-1* and *fca-3* mutants, as well as the parental Ler genotype, and assayed how individual cells varied in their *FLC* expression in these three genotypes at 7 days after sowing. We quantified the number of individual mRNAs per cell using single molecule Fluorescence *in situ* Hybridization (smFISH) (Fig. 2A,B, 2S1A-B). The endogenous Ler *FLC* carries a Mutator transposable element (TE) in intron 1, silencing *FLC* expression (Gazzani, Gendall et al. 2003, Michaels, He et al. 2003). Possibly because of the TE, we observed *FLC* mRNA accumulation in the nucleolus in this background (Fig. 2S1A). To avoid this complication, we transformed a Venus-tagged *FLC* into Ler and crossed that into our mutant genotypes (*fca-1*, *fca-3*) (Fig. 2A, (Berry, Hartley et al. 2015)). The transgenic *FLC* sequence was from Col-0, which does not contain the TE. We used smFISH probes for the Venus and *FLC* sequence. These probes generated a signal specific to the transgenic *FLC* copy in these plants, since we observed no signal in lines without the transgenic *FLC* (Fig. 2S1B). Comparison of whole-plant gene expression showed similar behaviour in the mutants for both the endogenous and transformed *FLC* (Fig. 1S1C,D). Focusing on the *Venus* sequence conferred the additional advantage that we were able to use the same lines for both mRNA and protein level quantifications (see below).

**Figure 2.**
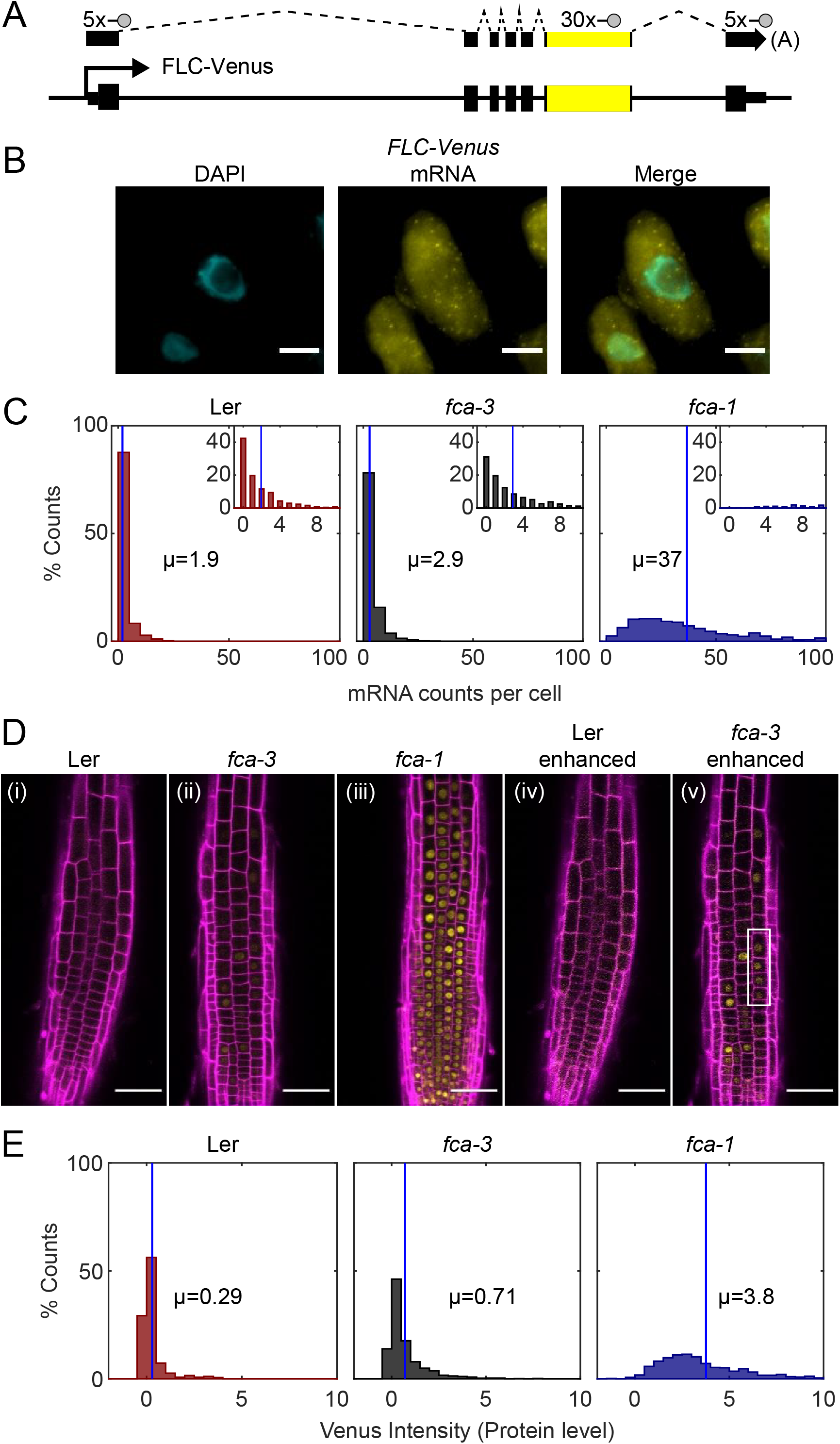
*FLC* expression per cell in *fca* mutants. **(A)** Schematic diagram of *FLC-Venus* locus with transcript and exonic probe position indicated. A total of 40 probes were designed, 10 against the *FLC* sequence and 30 for the *Venus* sequence. **(B)** Detection of *FLC-Venus* transcripts in single cells. Representative images of a cell with DAPI staining (cyan) and *FLC-Venus* mRNA (yellow) in *Arabidopsis thaliana* root cells. Scale bar, 5μm. **(C)** Histograms of smFISH results for each genotype (Ler, *fca-3*, *fca-1*) showing the number of single-molecule *FLC-Venus* RNAs detected per cell. The means of the distributions, µ, are indicated in each panel. Insets show the same data for mRNA counts between 0 and 10 (where 10 is the 95^th^ percentile of the *fca-3* data). For Ler n=853, *fca-3* n=1088, *fca-1* n=792; data from 7 days after sowing. Statistical tests: multiple comparisons following Kruskal-Wallis (*χ*^2^(2) = 1611.21, p-value = 0) with Tukey HSD post-hoc tests for *fca-3* – *fca-1*: p-value < 10^−8^; *fca-3* – Ler: p-value = 1.5 · 10^−5^; *fca-1* – Ler: p-value < 10^−8^. **(D)** Representative confocal images of roots for each genotype. Magenta shows the PI channel, Yellow is the FLC-Venus signal. Same settings were used for imaging and image presentation in (i)-(iii). Images in (iv) and (v) are the same as (ii) and (i), respectively, but adjusted to enhance the Venus signal by changing brightness and contrast. Scale bar, 50μm. **(E)** Histograms of FLC-Venus intensity per cell in each genotype. The means of the distributions, µ, are indicated in each panel. For Ler n=537 cells from 6 roots, *fca-3* n=1515 cells from 14 roots, *fca-1* n=1031 cells from 11 roots; data from 7 days after sowing. Statistical tests: multiple comparisons following Kruskal-Wallis (*χ*^2^(2) = 1607.56, p-value = 0) with Tukey HSD post-hoc tests for *fca-3* – *fca-1*: p-value < 10^−8^; *fca-3* – Ler: p-value < 10^−8^; *fca-1* – Ler: p-value < 10^−8^.

We observed that the average number of *FLC* mRNA transcripts per cell was highest in *fca-1*, lowest in Ler, and intermediate in *fca-3* (Fig. 2C, all differences are significant with *p* ≤ 1.5 · 10^−5^), with 89% of *fca-1* cells having higher expression than the 95th percentile of *fca-3* cells. Furthermore, when looking only at the ON cells (defined as cells with appreciable *FLC* expression, specifically more than one mRNA counted), the mRNA numbers in *fca-3* were only around 1/7 of those in *fca-1* (Fig. 2S1C), much less than one-half, ruling out that the reduced levels in *fca-3* were solely due to silencing of one of the two gene copies. Nevertheless, mRNA numbers in many *fca-3* cells were close to zero, suggesting that digital silencing may also be relevant in this case. Hence, the differences in overall expression between these three genotypes appeared to have two components. Firstly, there were different fractions of apparently silenced cells (digitally OFF) without any appreciable expression (one or no mRNAs counted): 62% in Ler, 51% in *fca-3* and 0.5% in *fca-1*. Secondly, in cells that did express *FLC* (digitally ON), the *FCA* mutations lead to an analog change in their expression levels, so that there was more *FLC* mRNA in *fca-1* ON cells than in the *fca-3* ON cells (Fig. 2S1C). This behaviour clearly differed from the digital regulation observed at *FLC* after cold treatment (Fig. 1A, (Angel, Song et al. 2011, Rosa, Duncan et al. 2016)).

We additionally used confocal live imaging to investigate FLC protein levels in individual cells (Fig. 2D, 2S2). After cell segmentation, we quantified the FLC-Venus intensity within the nuclei, comparing the different genotypes (Fig. 2E, 2S3A, all differences are significant with *p* < 10^−8^). This procedure allowed us to combine information of relative protein levels with the cell positions in the root. Mean intensity levels of FLC-Venus per cell and the overall histogram distribution, revealed again intermediate levels of FLC protein in *fca-3*, relative to Ler and *fca-1.* Mean levels in cells with appreciable protein amounts (ON cells) in *fca-3* were lower than half that in *fca-1* (Fig. 2D(i)-(iii), E, 2S3B), further supporting an analog regulatory component. At the same time, there was again strong evidence in favour of a digital component for *FLC* regulation (Fig. 2D(v)): in cells with the lowest protein levels, these levels were similar in *fca-3* and in Ler, again supporting a digital OFF state. Furthermore, we could see a mix of distinct ON and OFF cells by enhancing the *fca-3* images to increase the Venus intensity to a similar level as in *fca-1*. By contrast, in *fca-1* all cells were ON, whereas in Ler, cells appeared OFF even with an equivalent adjustment in this root (Fig. 2D(iv)), as did most other roots imaged, with only 15% of cells being ON in total (Fig. 2S3B). We could also infer a potentially heritable component in the ON/OFF states, as short files of ON cells could be observed in *fca-3* (Fig. 2D(v) white box). We additionally noticed a difference in *FLC* levels depending on the tissue, with *fca-3* epidermis cells having lower intensity on average than cells from the cortex (Fig. 2S3C). Overall, our results support a combination of analog and digital regulation for *FLC*: in Ler most cells were digitally OFF, in *fca-1* all cells were digitally ON, while in *fca-3* a fraction of cells were OFF, but for those cells that were ON, the level of *FLC* expression was reduced in an analog way relative to *fca-1*.

### *FLC* RNA and protein are degraded quickly relative to the cell cycle duration

Our results strongly pointed towards a digital switching component being important for *FLC* regulation by the autonomous pathway, but with an analog component too for those loci that remain ON. A study in yeast previously reported on the expected RNA distributions for the two cases of analog and digital control (Goodnight and Rine 2020). An important additional consideration when interpreting the analog/digital nature of the regulation concerns the half-lives of the mRNA and protein. In a digital scenario, long half-lives are expected to broaden histograms of mRNA/protein levels due to the extended times needed for mRNA/protein levels to increase/decrease after state switching. This could lead to a possible misinterpretation of analog regulation. In contrast, short half-lives will lead to clearer bimodality. Furthermore, what might look like a heritable transcription state could also appear due to slow dilution of a stable protein, as observed in other cases (Kueh, Champhekar et al. 2013, Zhao, Antoniou-Kourounioti et al. 2020). We therefore needed to measure the half-lives of the mRNA and protein to interpret our observations appropriately.

*FLC* mRNA has previously been shown to have a half-life of approximately 6 hours (Ietswaart, Rosa et al. 2017) in a different genotype (Col*FRI*) to that used here. We measured the half-lives of both RNA and protein in our highly expressing *FLC* line, *fca-1* (Fig. 3S1). The RNA half-life measurement used Actinomycin D treatment, inhibiting transcription, whereas the protein measurement used cycloheximide, arresting protein synthesis. The half-lives were then extracted from the subsequent decay in mRNA/protein levels. We found that both degradation rates were fast relative to the cell cycle duration, with half-lives of ∼5 and ∼1.5 hours respectively for the mRNA and protein (Fig. 3S1). Therefore, slow degradation is unlikely to be the cause of the apparent analog regulation and of the observed heritability seen in our root images. Furthermore, the short protein half-life indicates that any potential effects from growth causing dilution, and thus a reduction in protein concentrations, will also be small.

### One-way switching of *FLC* loci to an OFF state over time in *fca-3*

To understand the nature of potential digital switching, it is important to determine whether switching occurs from ON to OFF, OFF to ON, or in both directions. If all loci are switching one-way only, in either direction, this would lead to a gradual change of overall *FLC* expression over time. Alternatively, two-way switching or non-switching in at least a few cells would be necessary to have a constant concentration of *FLC* mRNA/protein over time. These considerations therefore raised the related question of whether cell population-level silencing is at steady state at the time of observation, or whether we are capturing a snapshot of a transient behaviour, with cells continuing to switch over developmental time.

To test if *FLC* expression is changing over time, we sampled *FLC* expression in the intermediate *fca-3* mutant, as well as *fca-1* and Ler, at 7, 15 and 21 days after sowing. This experiment revealed a decreasing trend in *fca-3* and Ler (Fig. 3A), which did not seem to be due to a change in *FCA* expression over the same timescale (Fig. 3B, compare particularly between 7 and 15 days). We therefore concluded that the most likely explanation was one-way switching from an otherwise heritable *FLC* ON state, to the heritable silenced OFF state occurring digitally and independently at individual loci in *fca-3* and Ler. In *fca-1* the trend was not statistically significant (*p* = 0.12). The potential absence of such a decrease in *fca-1* could be due to the high analog expression in *fca-1* preventing silencing in the majority of cells (see Discussion).

**Figure 3.**
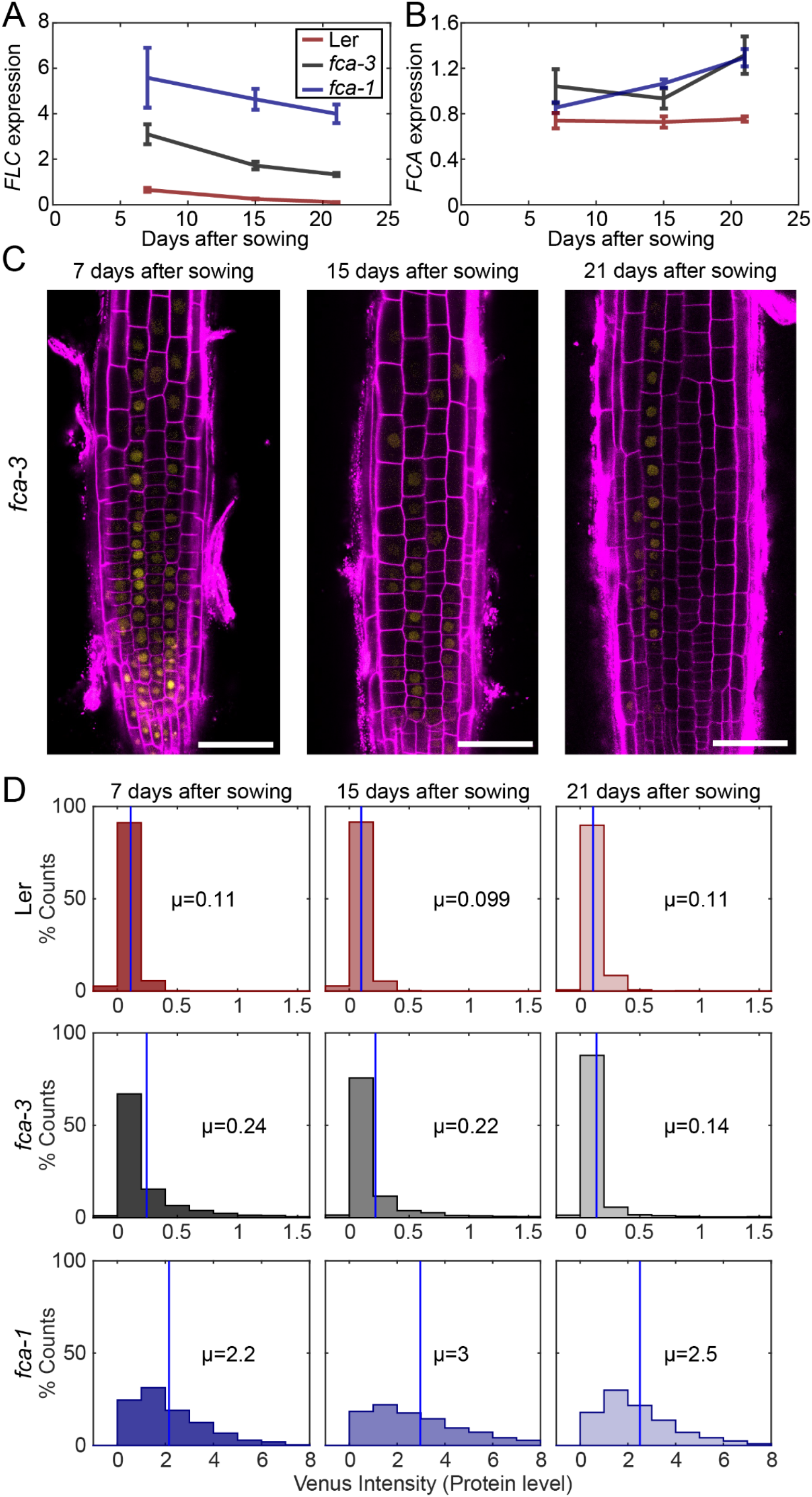
Experimental observation of gradual *FLC* silencing. **(A)** Timeseries of *FLC* expression in an *fca* mutant alleles transformed with the FLC-Venus construct. Expression is measured by qPCR relative to the house-keeping gene index (geometric mean of *PP2A* and *UBC)*. Error bars show SEM of n=3 biological replicates. Statistical tests: Linear regression on timeseries for each genotype. Slope for *fca-3* = −0.13, p-value = 2.0 · 10^−4^; Slope for *fca-1* = −0.11, p-value = 0.12; Slope for Ler = −0.038, p-value = 4.8 · 10^−6^. **(B)** Timeseries of *FCA* expression, otherwise as in (A). **(C)** Representative images of *fca-3* roots by confocal microscopy. Magenta shows the PI channel, Yellow is the FLC-Venus signal. Same settings were used for imaging and image presentation. Scale bar, 50 μm. **(D)** Histograms of FLC-Venus intensity per cell in *fca-3* at each timepoint. The means, µ, of the distributions are indicated in each panel. For Ler: 7 days, n=1121 cells from 10 roots; 15 days, n=1311 cells from 9 roots; 21 days, n=1679 cells from 12 roots. For *fca-3*: 7 days, n=2875 cells from 24 roots; 15 days, n=3553 cells from 23 roots; 21 days, n=3663 cells from 21 roots. For *fca-1*: 7 days, n=1022 cells from 9 roots; 15 days, n=1770 cells from 12 roots; 21 days, n=2124 cells from 12 roots. Statistical tests: Linear regression on timeseries for each genotype. Slope for *fca-3* = −0.0077, p-value = 4.0 · 10^−46,^; Slope for *fca-1* = 0.018, p-value = 0.00064; Slope for Ler = −0.00015, p-value = 0.44.

By imaging FLC-Venus we observed that the fraction of ON cells was indeed decreasing in *fca-3*, over the time course (Fig. 3C,D, 3S2). In fact, after 21 days the pattern of ON/OFF cells in the *fca-3* roots was very similar to that of plants that had experienced cold leading to partial cell population-level *FLC* shutdown, with the majority of cell files being stably repressed, but still with some long files of cells in which *FLC* was ON (compare timepoint 21 in Fig. 3C and figures in (Berry, Hartley et al. 2015)). It remains possible that there is a low level of switching in the opposite direction, from OFF to ON, in *fca-3*. The other genotypes did not show this statistically significant decreasing trend in the FLC-Venus data. In *fca-1*, the fraction of ON cells did not show a consistent pattern, while the qPCR data showed a non-statistically significant decrease (Fig. 3A), indicating an overall lack of switching. In Ler, the fraction of ON cells remained essentially constant (at a very low level), differing from the qPCR data, where a statistically significant reduction was observed (Fig. 3A). This result suggests that root tip cells in Ler could switch off early, while ON cells still remain at the whole plant level that continue to switch off, thereby explaining the decrease in the qPCR experiment (Fig. 3A).

We also examined whether the change over time in *fca-3* could be due to an analog change in the expression of the ON cells rather than a decreasing number of ON cells. By setting a fixed threshold at 0.5 intensity, separating ON and OFF cells, we found that the histogram of *fca-3* cells with intensities above that threshold (normalised by the total number of these cells in each condition) (Fig. 3S3) was similar at all timepoints. This finding is consistent with digital regulation: a decrease in the number of ON cells, but with all remaining ON cells expressing *FLC* at similarly high levels over time. Overall, our results are consistent with a progressive one-way digital switch to OFF for *FLC* loci in *fca-3*.

### Mathematical model incorporating one-way switching can recapitulate the *FLC* distribution over time in *fca-3*

We finally developed a mathematical model for *FLC* RNA/protein levels in the root (Fig. 4A, 4S1, Materials and Methods), to test if the analog and digital components inferred from the data were sufficient to reproduce the experimentally observed patterns. The model incorporated digital *FLC* state switching embedded in simulated dividing cells in the root. Construction of the model took account of many of the above results on *FLC* dynamics, including rapid turnover of the RNA and protein (giving sharp switching) and directionality of the switching (ON to OFF). We focused on modelling the ON to OFF switching dynamic and used distributions for the protein levels in cells with two ON loci, one ON and one OFF, and two OFF empirically fitted to our data (Fig. 4A, Materials and Methods). These empirical distributions will include any possible effects from cell size and burstiness.

**Figure 4.**
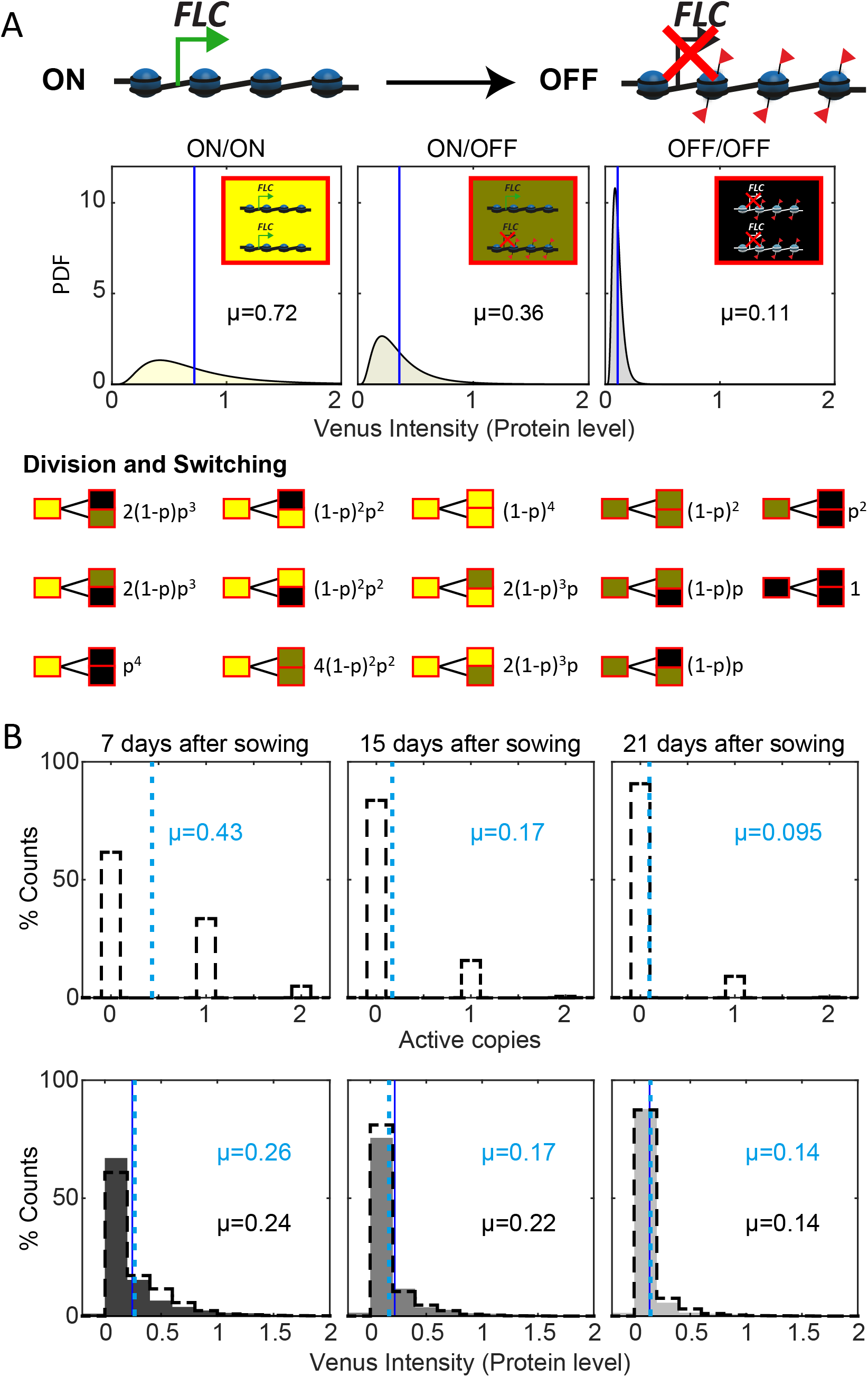
Mathematical model captures *FLC* regulation. **(A)** Diagram of mathematical model. Individual *FLC* gene copies can be ON or OFF, such that a cell can be in one of three states depending on the combination of ON and OFF gene copies within it (ON/ON, ON/OFF, OFF/OFF). Venus intensity (corresponding to amount of protein) within a cell was sampled from the distributions shown (described in Methods section), depending on the cell state. The means of the distributions, µ, are indicated in each panel. Switching of ON *FLC* gene copies to OFF can occur at division (with probability *p* = 0.25 per division): neither, one or both copies can switch OFF in each daughter cell at each division. **(B)** Histograms of active *FLC* copies per cell (top) and Venus intensity per cell (bottom) at the indicated timepoints. Model histograms are plotted with dashed lines around empty bars and experimental data is shown as filled grey histograms with no outline. The means of the distributions, µ, are indicated in each panel for the data (black text and dark blue solid line) and the model (light blue text and dashed line). Model simulated 500 *fca-3* cell files and histograms shown exclude bottom 4 cells of each file, as cells near the QC are also not included in the imaging.

Another potentially confounding issue is that in the plant root there is a mix of dividing and differentiated cells. Our experimental observations capture cells primarily in the division zone of the root, but even within this region, cell cycle times are not the same in all cells (Rahni and Birnbaum 2019). Incorporating this heterogeneity is important as digital chromatin inheritance is affected by DNA replication (Baxter, Šviković et al. 2021) and therefore switching is likely to be different between dividing and differentiated cells, and to occur during/after replication. In our model, we assumed that stochastic ON to OFF switching occurred during cell division to capture this effect of the cell cycle, and we used cell cycle lengths based on the literature (Rahni and Birnbaum 2019).

With these assumptions, our model could be fit (Materials and Methods) to replicate the observed pattern of increasing OFF cells in *fca-3* roots, as well as the quantitative histograms for protein levels in *fca-3* (Fig. 4B). In addition, the model could also capture the longer files present in the later timepoints in the data (T21, Fig. 3C, 4S1), although the prevalence of this effect in the data may suggest additional developmental influence on the heritability of the ON/OFF state at later times. Overall, however, the model incorporating one way ON to OFF switching can faithfully recapitulate the developmental dynamics of *FLC* in *fca-3*.

## Discussion

In this work we have uncovered a combination of analog and digital transcriptional regulation for the gene *FLC*: analog regulation arises through the autonomous pathway, as illustrated by the *fca* mutants, before digital switching into a Polycomb silenced state. In the case of *fca-3* with intermediate *FLC* expression, this digital switching occurs slowly and so can be observed in our time series experiment, underlining a clear temporal separation of analog and digital transcriptional control. However, for Ler, the switch appears to mostly occur already before our first experimental measurements at 7 days, whereas in *fca-1* the switch does not occur at all.

The analog autonomous pathway component of *FLC* regulation controls transcription through an FCA-mediated mechanism. Based on previous studies on the interplay between transcription and PRC2 silencing (Beltran, Yates et al. 2016, Berry, Dean et al. 2017, Holoch, Wassef et al. 2021, Lövkvist, Mikulski et al. 2021), we propose that digital Polycomb silencing is prevented by high transcription in *fca-1*, while low transcription is not sufficient to significantly oppose silencing in Ler. In *fca-3*, the dynamical antagonism between Polycomb silencing and an intermediate level of transcription is still ongoing at our experimentally accessible timepoints. In this mutant, an initial transcriptionally active ON state with intermediate transcription levels controlled by an analog mode of autonomous pathway regulation is eventually overcome and stochastically switched at each gene copy into a digital Polycomb silenced OFF state. In this way, both analog and digital regulation are combined at *FLC*, with the timescale for one-way switching between these modes controlled by the strength of initial analog transcription. In *fca-3* such switching to the OFF state causes a gradual reduction in *FLC* expression at the whole plant level. Interestingly, we found that the mRNA levels/FLC-Venus intensities of ON cells in Ler and *fca-3* are similar (Fig. 2S1C, 2S3B). This result could be due to a more rapid switch-off of loci which happen to have lower levels of transcription, leaving only more highly expressing cells that are able to stay ON, even in Ler. Furthermore, we emphasise that there may be additional developmentally regulated processes occurring at *FLC* in addition to a simple one-way OFF switch with a constant probability in each genotype. This possibility is underscored by the model not completely recapitulating the long cell files we observe experimentally at the 21-day timepoint.

A prolonged environmental cold signal can also result in a switch from cells expressing *FLC* to non-expressing OFF cells, a process which occurs independently at each gene copy of *FLC* (Berry, Hartley et al. 2015). These ON/OFF states are then maintained through cell divisions leading to epigenetic memory. Here, we have addressed how *FLC* is regulated in the absence of cold and we show that a similar mode of regulation occurs. Furthermore, at least in the case of *fca-3* with initially intermediate transcription levels, silencing at the cell population-level also happens gradually over many cell cycles, similar to the behaviour over prolonged periods of cold. Moreover, images of roots at 21 days after sowing without cold, with fluorescently labelled FLC, are visually strikingly similar to images of vernalized roots, where in both cases we find whole files of either all ON or all OFF cells. These comparisons suggest similarities in the molecular characteristics of the Polycomb silenced state at *FLC* generated either with or without cold treatment.

The main antagonist of the autonomous pathway is the *FRIGIDA* gene, a transcriptional activator of *FLC* (Fig. 1B), which was not present in the lines used in this study. For future work, it would be interesting to investigate transcriptional responses in a *FRI* allelic series. This could be particularly important because natural accessions show mutations in *FRI* rather than in autonomous pathway components, possibly because the autonomous components regulate many more target genes.

Overall, our work has revealed a combination of both analog and digital modes of regulation at Arabidopsis *FLC* before cold, with analog preceding digital. We further propose it is the strength of the initial analog, autonomous pathway transcriptional level that controls the timescale for the switch into the subsequent digital, Polycomb silenced state.

## Materials and Methods

### Plant growth

Seeds were surface sterilized in 5% v/v sodium hypochlorite for 5 min and rinsed three times in sterile distilled water. Seeds were stratified for 2 days at 4°C in petri dishes containing MS media without glucose. The plates were placed vertically in a growth cabinet (16 hours light, 22°C) for 1 week.

### Gene expression analysis

For gene expression time series, 20+ plants were harvested at each timepoint (7, 15, 21 days old plants). Plant material was snap frozen with liquid nitrogen, ground and RNA was extracted using the Phenol:Chloroform:Isoamyl alcohol (25:24:1) protocol (Yang, Howard et al. 2014). RNA was purified with the TurboDNase (Ambion) kit to remove DNA contamination and reverse transcribed into cDNA using SuperScript IV Reverse transcriptase (Invitrogen) and RT primers for genes of interest. Gene expression was measured by qPCR, data was normalized to PP2A and UBC. Primer sequences are summarized in Supplementary Table 1.

### Actinomycin D treatment

For Actinomycin D (ActD) experiments, 6-day-old plants were initially germinated in non-supplemented media and were transferred to new plates containing ActD. Stock solution of ActD (1mg/mL dissolved in DMSO) was added to molten MS media to a final concentration of 20μg/mL. ActD was obtained from SigmaTM (catalogue # A4262-2MG).

### Cycloheximide (CHX) treatment

FLC-Venus protein stability was assayed with the de novo protein synthesis inhibitor cycloheximide (C1988, Sigma-Aldrich) following the procedures in (Zhao, Antoniou-Kourounioti et al. 2020). Briefly, 7-day old *fca-1* seedlings carrying the *FLC-Venus* transgene were treated in liquid MS medium containing 100μM CHX. The seedlings were sampled after 0, 1.5, 3, 6, 12 and 24 hours of treatment. A non-treatment control is also included, in which seedlings were soaked in liquid MS medium without the inhibitor CHX for 24 hours. Approximately 1.0ng seedlings were ground to a fine powder with Geno/Grinder. Total protein was extracted with 1.5 mL buffer (50 mM Tris-Cl pH 8.0, 15 4mM NaCl, 5 mM MgCl_2_, 10% glycerol, 0.3% NP-40, 1% Triton-100, 5 mM DTT and protease inhibitor). Then each sample was cleaned by centrifugation at 16,000g at 4°C for 15 min, 50 μl total protein was taken as input before enrichment with magnetic GFP-trap beads (GTMA-20, Chromo Tek). The input samples were run on a separate gel and used as a processing loading control for the starting level for each sample. The enriched FLC-Venus protein was detected by western blot assay with the antibody anti-GFP (11814460001, Roche). Signals were visualized with chemiluminescence (34095, Pierce) with a secondary antibody conjugated to Horseradish peroxidase (NXA931V, GE Healthcare). The chemiluminescence signal was obtained by the FUJI Medical X-ray film (4741019289, FUJI). Quantification was performed with ImageJ after the films were scanned with a printer scanner (RICOH). Ponceau staining was performed with commercial Ponceau buffer (P7170, Sigma-Aldrich) and used as the processing controls. All of the western blot assays were performed with equal weight of whole seedlings.

### smFISH on root squashes

smFISH was carried out as described by Duncan et al. 2016. Briefly, root tips from 7-day-old seedlings were cut using a razor blade and placed into glass wells containing 4% paraformaldehyde and fixed for 30 min. Roots were then removed from the fixative and washed twice with nuclease free 1X PBS. Several roots were then arranged on a microscope slide and squashed between the slide and coverslip. Slides were submerged (together with the coverslips) for a few seconds in liquid nitrogen until frozen. The coverslips were then removed, and the roots were left to dry at room temperature for 30 min.

Tissue permeabilization and clearing were achieved by immersing sequentially the samples in 100% methanol for 1h, 100% ethanol for 1h and 70% ethanol for a minimum of one hour. The ethanol was left to evaporate at room temperature for 5 min and slides were then washed with Stellaris RNA FISH Wash Buffer A (Biosearch Technologies Cat# SMF-WA1-60). 100 μL of hybridization solution (containing 10% dextran sulfate, 2x SSC and 10% formamide), with each probe set at a final concentration of 125 nM, was then added to each slide. The slides were left to hybridize at 37°C overnight in the dark.

The hybridization solution containing unbound probes was pipetted out the following morning. Each sample was then washed twice with Stellaris RNA FISH Wash Buffer B (Biosearch Technologies Cat# SMF-WB1-20) with the second wash left to incubate for 30 min at 37°C. 100 μl of the nuclear stain DAPI (100 ng/mL) was then added to each slide and left to incubate at 37°C for 10 minutes. Slides were then quickly washed with 2xSSC. 100 μl GLOX buffer minus enzymes (0.4% glucose in 10 mM Tris, 2X SSC) was added to the slides and left to equilibrate for 2 min. Finally, this was removed and replaced with 100 μL of GLOX buffer (containing 1 μl of each of the enzymes glucose oxidase (#G0543 from Sigma) and catalase (#C3155 from Sigma)). The samples were then covered by 22 mm × 22 mm No.1,5 coverslips (VWR), sealed with nail varnish and immediately imaged.

### smFISH Probe synthesis

We used the online program Stellaris Probe Designer version 2.0 from Biosearch Technologies to design probe sequences for *FLC-Venus* and unspliced *PP2A*. For probe sequences, see Supplementary Tables 2 and 3.

### Image Acquisition

The smFISH slides were imaged using a Zeiss LSM800 inverted microscope, with a 63x water-immersion objective (1.20 NA) and Microscopy Camera Axiocam 503 mono. The following wavelengths were used for fluorescence detection: for probes labelled with Quasar570 an excitation filter 533-558 nm was used and signal was detected at 570-640 nm; for probes labelled with Quasar670 an excitation filter 625-655 nm was used and signal was detected at 665-715 nm; for DAPI an excitation filter 335-383 nm was used and signal was detected at 420-470 nm; for GFP an excitation filter 450-490 nm was used and signal was detected at 500-550 nm.

For the FLC-Venus protein level quantification, optical sections of roots were collected with a Zeiss LSM780 microscope equipped with a Channel Spectral GaAsP detector with a 20x objective (0.8 NA). For z-stacks, the step size was 0.5 μm with a pinhole aperture of 1.5 AU. The overall Z size varied between 45 to 75 slices depending on the orientation of the root. Roots from FLC-Venus lines were immersed in 1 μg/mL propidium iodide (PI, Sigma–Aldrich, P4864) to label the cell wall. For visualization of roots stained with propidium iodide, an excitation line of 514 nm was used and signal was detected at wavelengths of 611 to 656 nm. For observation of Venus signal, we used a 514 nm excitation line and detected from 518 to 535 nm. To allow comparison between treatments, the same laser power and detector settings were used for all FLC-Venus images.

In FLC-Venus time course imaging of epidermal root meristems, the Leica SP8X or Stellaris 8 were used with 20x multi-immersion objective (0.75 NA). The Argon (SP8X) or OPSL 514 (Stellaris 8) lasers were used at 5% to excite FLC-Venus and PI at a 514 nm wavelength in bidirectional mode (PI signal was used for set-up). Venus was detected between 518-550 nm with the HyD SMD2 detector in photon counting mode; PI was detected at 600-675 nm. Epidermal images were obtained in photon-counting mode with laser speed 200, line accumulation of 6 (pixel dwell time of 2.43 µs) and a Z-step size of 0.95 µm and a pinhole size of 1 AU.

For representative images, these were projected such that one single middle slice from the PI channel was used to show the cell outline, onto which 10 slices of FLC-Venus channel were average intensity projected (T7, LSM780 imaging, Fig. 2) or sum projected for time-course imaging (T7/T15/T21, SP8X imaging, Fig. 3). The dynamic range of the FLC-Venus signal was pushed from 0-255 to 4-22 (for LSM780 images, Fig.2) and 0-20 (for SP8X images, Fig. 3) for all images apart from where enhanced. In enhanced *fca-3*/Ler images, the dynamic range was further pushed to 5-10 (for LSM780 images, Fig.2) and to 0-6 (for SP8X images, Fig. 3) to obtain a similarly strong signal as observed in *fca-1*.

### Image analysis

#### FISH analysis

Quantification of FISH probes took place in two stages:

1. Identification of probe locations in the whole 3D image, excluding the top and bottom z-slice from each z-stack due to light reflection at the plant cell wall.
2. Assignment of identified probes to specific cells via segmentation of the image into regions.

To detect probes, a white tophat filter was applied to the probe channel, followed by image normalisation and thresholding. Individual probes were then identified by connected component segmentation. The centroids of each segmented region were assigned as the probe’s location.

Images were manually annotated with markers to indicate positions of nuclei and whether cells were fully visible or occluded. A combination of the visibility markers and nuclei were used to seed a watershed segmentation of the image, thus dividing the image into regions representing individual cells. Probes within each region were counted to generate the per-cell counts. Occluded cells were excluded from the analysis (probe counts for those regions were ignored).

Custom code is available from: https://github.com/JIC-Image-Analysis/fishtools.

### FLC-Venus fluorescence intensity

To measure per-cell FLC-Venus intensities, we developed a custom image analysis pipeline to extract cell structure information from the PI cell well information and use this to measure per-cell nuclear fluorescence. Images were initially segmented using the SimpleITK (Beare and Lehmann 2006) implementation of the morphological watershed algorithm. Reconstructed regions touching the image boundaries and those below a certain size threshold were removed. Segmentations were then manually curated to merge over-segmented regions, and to assign file identities to the resulting segmented cells. This curation was performed using custom software, able to merge segmented cells, resegment cells from specified seeds, and split cells along user-defined planes. To approximate the nuclear position, we fitted a fixed size spherical volume of 15 voxels radius to the point of maximal FLC-Venus intensity in each reconstructed cell. Per-voxel mean FLC-Venus intensities were then calculated by dividing the summed intensity within the spherical region by the sphere volume.

Custom code for initial segmentation and Venus intensity measurement is written in the Python language (Python), and is available at https://github.com/JIC-Image-Analysis/root_measurement.

The code for the segmentation curation software is available at https://github.com/jfozard/segcorrect.

### Mathematical model

We constructed a model for *FLC* chromatin states and protein levels in the root (Fig. 4A) and used it to simulate root cell files and generate simulated protein level histograms (Fig. 4B, 4S1).

The root was represented as a collection of cell files (such as the 12 representative cell files shown in Fig. 4S1), and 500 cell files were simulated in total. Each cell file consisted of a list of 30 cells (Supplementary Table 4), ordered from the root tip toward the rest of the plant. This region was intended to approximately match the division zone and imaging region. Cells were able to divide, giving rise to two daughter cell and pushing cells that were further from the tip upwards. Cells escaping the 30-cell limit were removed from the simulation. Cells in a given file were clonally related to each other, and eventually all originated from divisions of the “initial cells” (the first cells of the cell file, which are adjacent to the quiescent centre (QC) of the root apical meristem). Each cell was described by its index along the cell file, its digital chromatin state, its protein concentration (expressed as the Venus intensity to match experimental observations) and its remaining cell cycle duration.

With regards to the digital chromatin state, a cell could be in an ON/ON (both *FLC* copies in the active chromatin state), ON/OFF (one active and one inactive) or OFF/OFF state (both inactive). All cells at the start of the simulation at 0 days are in the ON/ON configuration. The model also included a 1-way switching process from a heritable ON state to a heritable OFF state that occurs in a cell-cycle dependent manner (Baxter, Šviković et al. 2021). The assumption of one-way switching from ON to OFF was motivated by our data (Fig. 3), as well as the fact that wildtype Ler loci are primarily OFF, and therefore the presence of ON cells in the *fca* mutants might be explained by a disruption of the OFF switch. A single *FLC* copy could switch from an ON to an OFF state during division with a probability *p* = 0.25 (Supplementary Table 4). The possible transitions of the combined two copies in a cell, and their probabilities, are shown in Fig. 4A. A natural consequence of one-way switching is that, over time, the *FLC* levels decrease in the simulated root for *fca-3* as more and more loci switch to OFF (Fig. 4B), as was observed experimentally (Fig. 3).

In order to compare the simulated cell states against the FLC-Venus data, we processed the model outputs using an additional step. This step gave each ON *FLC* copy an associated protein level, which was sampled from a log-normal distribution with parameters: 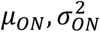. The background within each cell was sampled from a log-normal distribution with parameters: 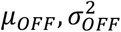. These four parameters and the switch-off probability *p* were manually fitted to the experimental Venus distributions for *fca-3* at 7, 15 and 21 days (Fig. 3D, 4, Supplementary Table 4). Protein levels for each cell, according to the combination of ON and OFF *FLC* copies (Fig. 4A), were given by appropriate combinations of random variables, specifically,

for ON/ON cells: 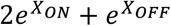
for ON/OFF cells: 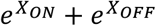
and for OFF/OFF cells: 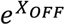

where 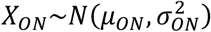 and 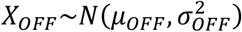. The noisy nucleus background signal (*X_OFF_*), was assumed to be higher than any real signal originating from OFF *FLC* copies.

The duration of the cell cycle of each cell was determined by the cell’s position along the cell file at the time of the division that created it. These cell cycle times according to position were based on the literature (Rahni and Birnbaum 2019). We used a truncated normal distribution with the mean and standard deviation matching the measured values for epidermis cells. A minimum value was set such that the cell cycle could not be shorter than 13 hours (the lowest value observed for epidermis or cortex cells in (Rahni and Birnbaum 2019)).

Custom code is available from: https://github.com/ReaAntKour/fca_alleles_root_model.

## Acknowledgements

We thank all members of the Dean, Rosa and Howard groups for excellent discussions. Special thanks to Dr. Silvia Costa for imaging advice and Dr. Cecilia Lövkvist for project coordination. This work was supported by BBSRC Institute Strategic Programme GEN (BB/P013511/1), BBSRC grant BB/P020380/1 to C.D. and M.H., European Union’s Horizon 2020 research and innovation programme (Marie Skłodowska-Curie grant No. 813282) to Sv.R. and M.H. and Vetenskapsrådet grant 2018-04101 to A.M. and S.R.

## Author Contributions

MH, CD, SR, RAK, AM, SvR, GM, SB designed research, AM, SvR, SB, YZ, HW performed research, MHa built image analysis pipeline with modifications by RAK, JF built python script for cell segmentations, RAK, SvR, AM, TH analysed data, RAK built mathematical model, RAK, AM, SvR and MH wrote the manuscript with help from all authors. All authors agreed on the final version.

## Code availability

All code is available through github repositories, as described in the Materials and Methods.

## Data availability

Image data used in this study are available in the BioImage Archive under accession number S-BIAD425 (https://www.ebi.ac.uk/biostudies/studies/S-BIAD425). Derived quantification data from images, as well as qPCR data, are provided as Supplementary file 1.

---

**Figure 1 - Supplementary 1.**
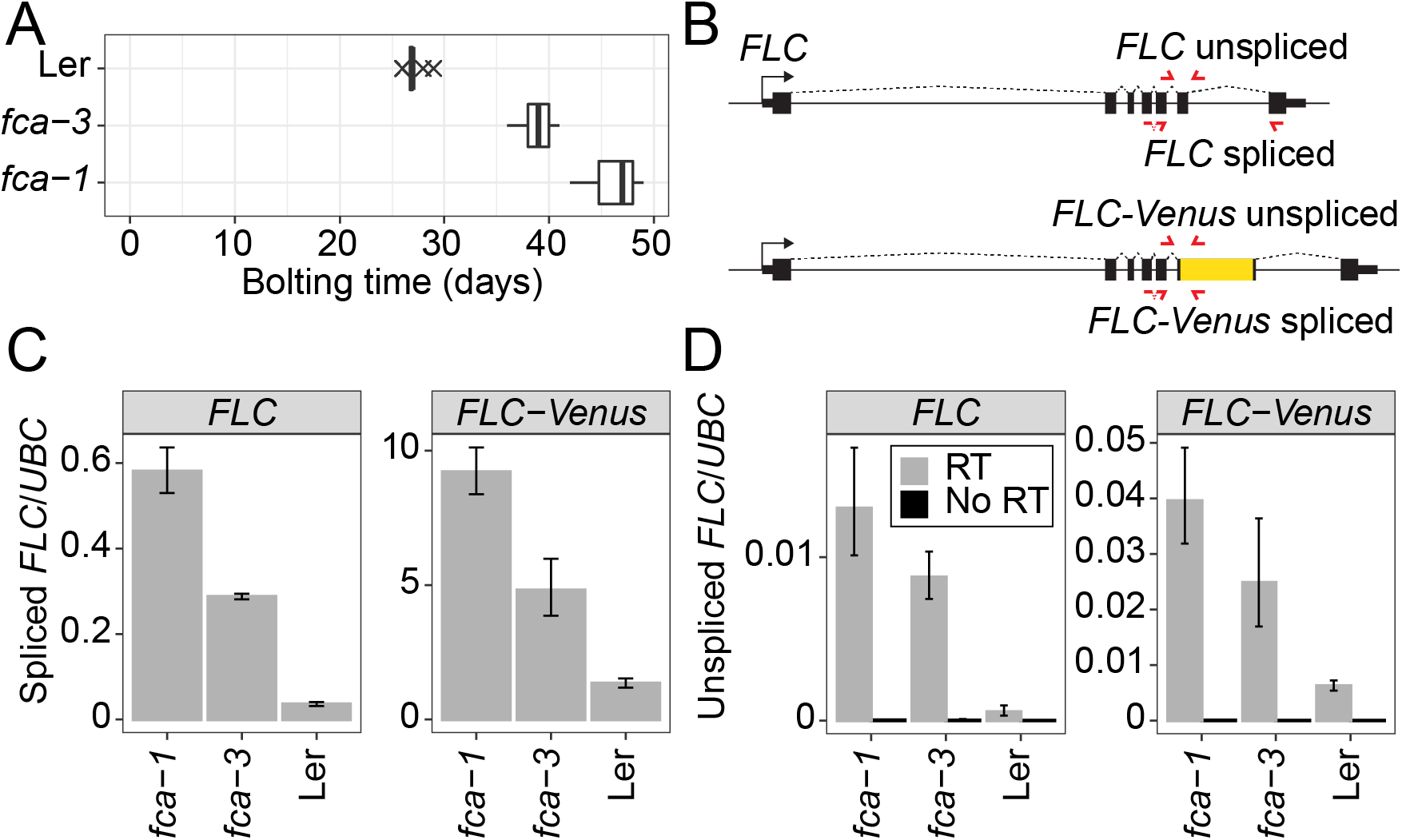
Characterisation of *fca* alleles and FLC-Venus transgene. **(A)** Flowering time measured in days from sowing until first appearance of a floral meristem (bolting). 12 plants were measured for each genotype. Boxplots show median and interquartile range (IQR), with data points outside median ± 1.5 IQR shown as crosses. **(B)** Diagram showing positions of qPCR primers (red) to quantify spliced and unspliced endogenous *FLC* and transgenic *FLC-Venus* RNA. **(C-D)** *FLC* and *FLC-Venus* RNA measured by qPCR, relative to *UBC* RNA. No RT (Reverse Transcription) negative control in D shows no expression with same unspliced primers. Error bars show SEM of n≥2 biological replicates, harvested 14 days after sowing.

**Figure 2 - Supplementary 1.**
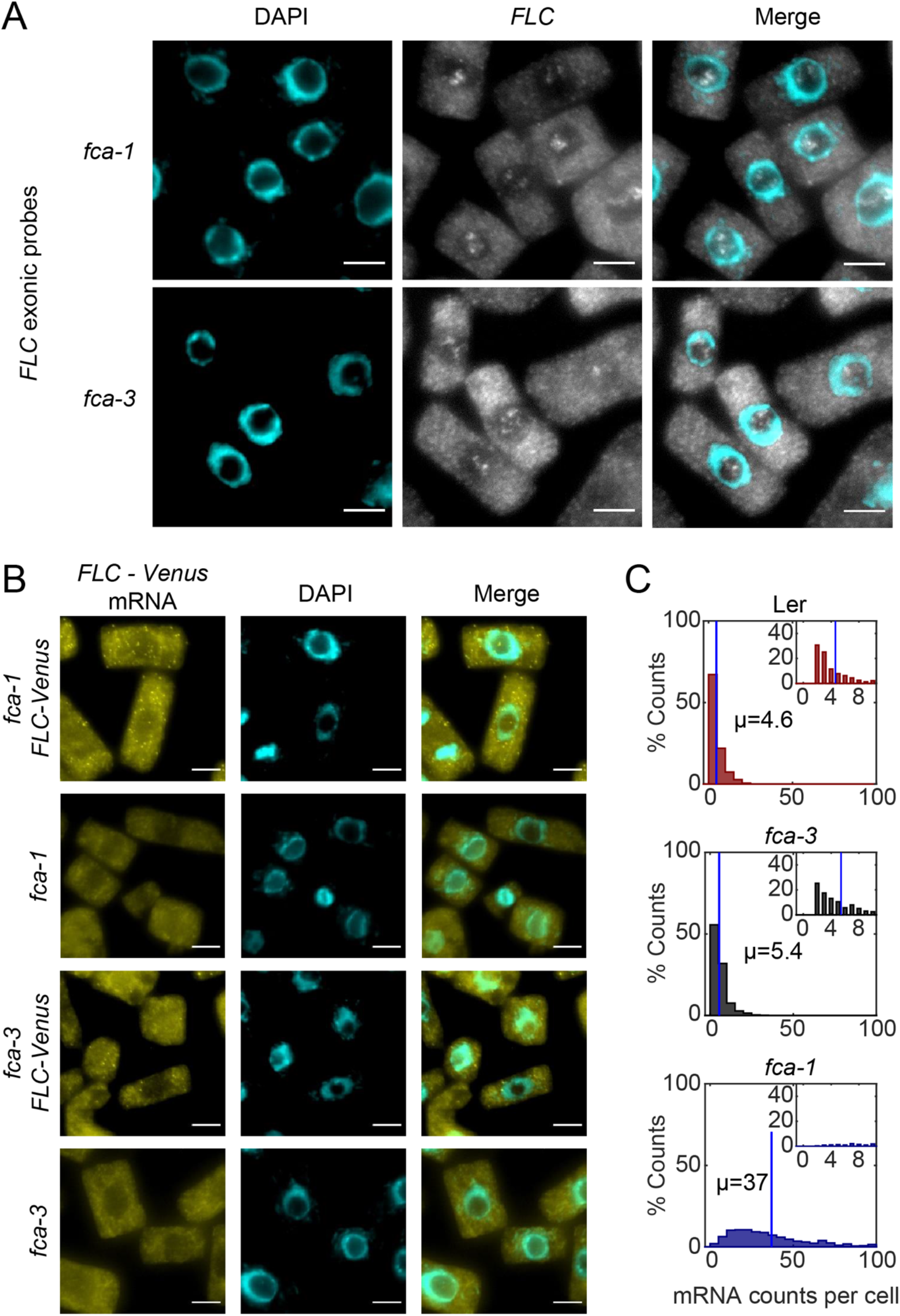
smFISH method for FLC-Venus imaging. **(A)** Detection of nucleolar signal using *FLC* specific probes in Ler background *fca-1* and *fca-3* plants in *Arabidopsis thaliana* root cells. The nucleus is stained with DAPI (cyan). Scale bar, 5 μm. **(B)** FLC-Venus probes bind specifically to the transgenic *FLC-Venus* RNA and not to the endogenous *FLC* transcript. All data from seedlings 7 days after sowing. Scale bar, 5 μm. **(C)** The data of Fig. 2C plotted for the three genotypes but showing only measurements in “ON cells”. These are defined as cells with more than one *FLC-Venus* mRNA. The means of the distributions, µ, are indicated in each panel. For Ler n=323, *fca-3* n=536, *fca-1* n=788, from 7 days post sowing. Statistical tests: multiple comparisons following Kruskal-Wallis (*χ*^2^(2) = 1067.37, p-value = 1.7 · 10^−232^) with Tukey HSD post-hoc tests for *fca-3* – *fca-1*: p-value < 10^−8^; *fca-3* – Ler: p-value = 0.12; *fca-1* – Ler: p-value < 10^−8^.

**Figure 2 - Supplementary 2.**
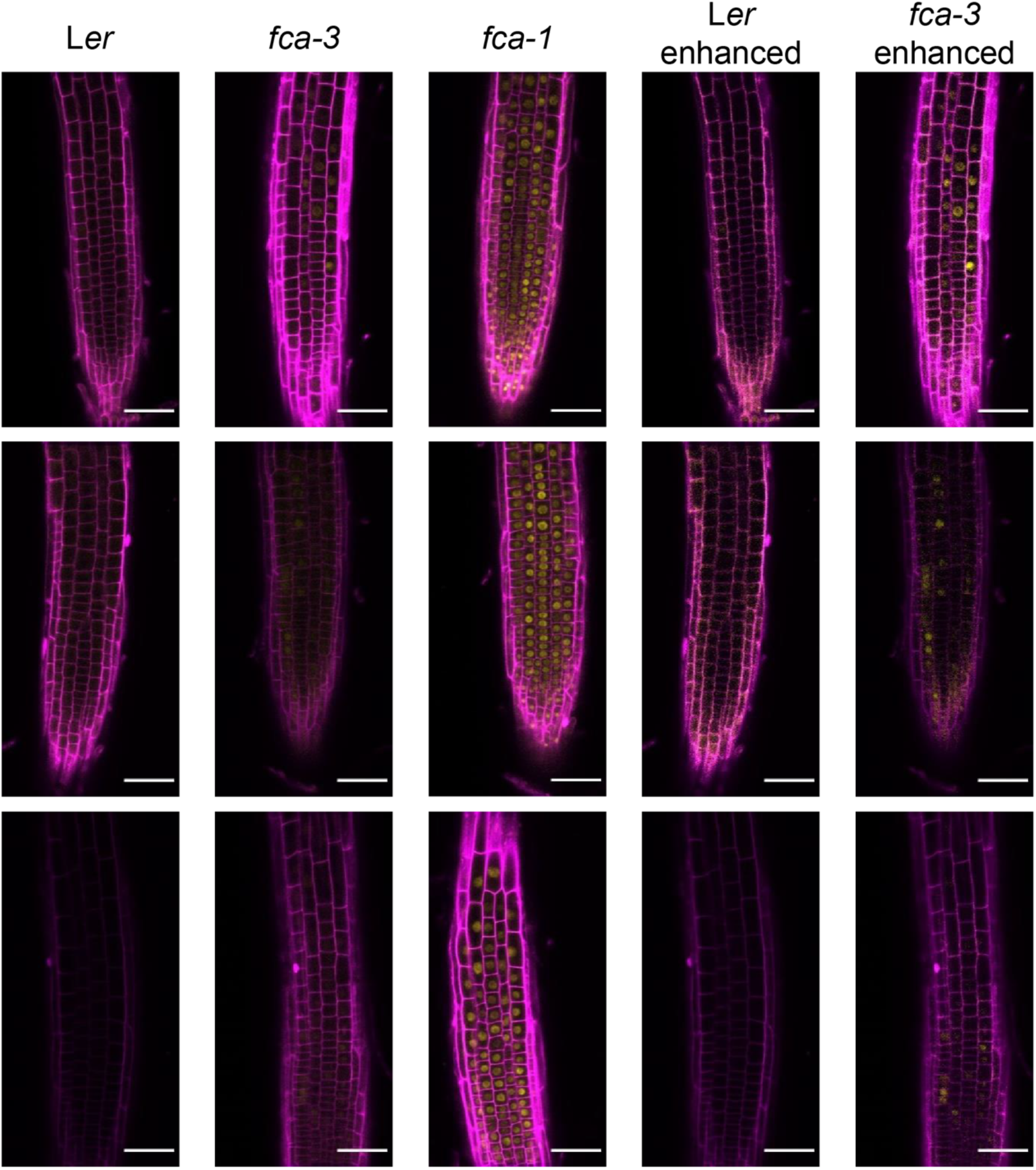
FLC-Venus imaging in *fca* alleles – root replicates. Additional representative images of FLC-Venus signal in the three genotypes (Ler, *fca-3*, *fca-1*) from 7 days post sowing. Magenta shows the PI channel, Yellow is the FLC-Venus signal. Same settings were used for imaging and image presentation in left three columns. “Enhanced” images (two columns on right) are the same respective images, adjusted to enhance Venus signal by changing brightness and contrast. Scale bar, 50 μm.

**Figure 2 - Supplementary 3.**
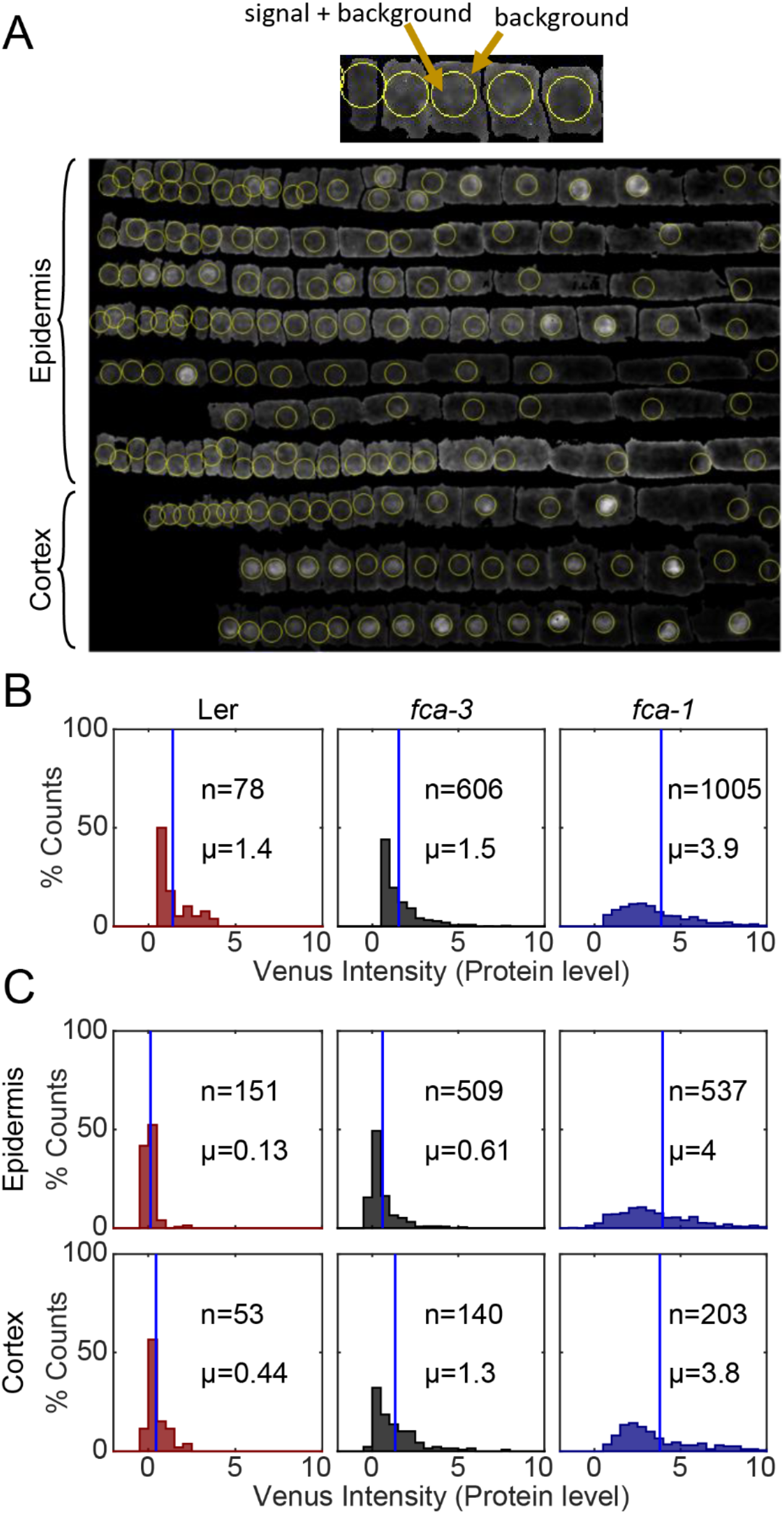
Segmentation and quantification method. **(A)** Custom visualisation and segmentation software was used to manually correct automatic segmentations and annotate cells according to cell file. Top: Diagram explaining quantification of FLC-Venus intensity. In ON cells, the sphere contains the nucleus and therefore the FLC-Venus signal as well as the background signal. In OFF cells, the sphere will contain only background signal. The mean intensity outside the sphere was used as the background signal and this was subtracted from the sphere intensity. Bottom: Cell files from a single root separated according to tissue type demonstrating the result of the nucleus finding algorithm. Longitudinal section shown. In each cell, the sphere of constant radius that maximised fluorescent intensity inside the sphere was determined. For each cell, the position of the sphere is shown, corresponding to the nucleus in ON cells. **(B)** The data of Fig. 2E plotted for the three genotypes but showing only measurements in “ON cells”. These are defined as cells with intensity higher than 0.5. The means of the distributions, µ, and number of cells, n, are indicated in each panel. Statistical tests: multiple comparisons following Kruskal-Wallis (*χ*^2^(2) = 552.88, p-value = 8.8 · 10^−121^) with Tukey HSD post-hoc tests for *fca-3* – *fca-1*: p-value < 10^−8^; *fca-3* – Ler: p-value = 0.57; *fca-1* – Ler: p-value < 10^−8^. **(C)** Cortex files are more likely to be ON and/or have higher intensity. Histograms showing FLC-Venus intensity in cells separated according to tissue type; data from seedlings 7 days post sowing. The means of the distributions, µ, and number of cells, n, are indicated in each panel. Statistical tests: Two-sample t-test (t-statistic: −7.66, df: 647, p-value = 6.7 · 10^−14^).

**Figure 3 - Supplementary 1.**
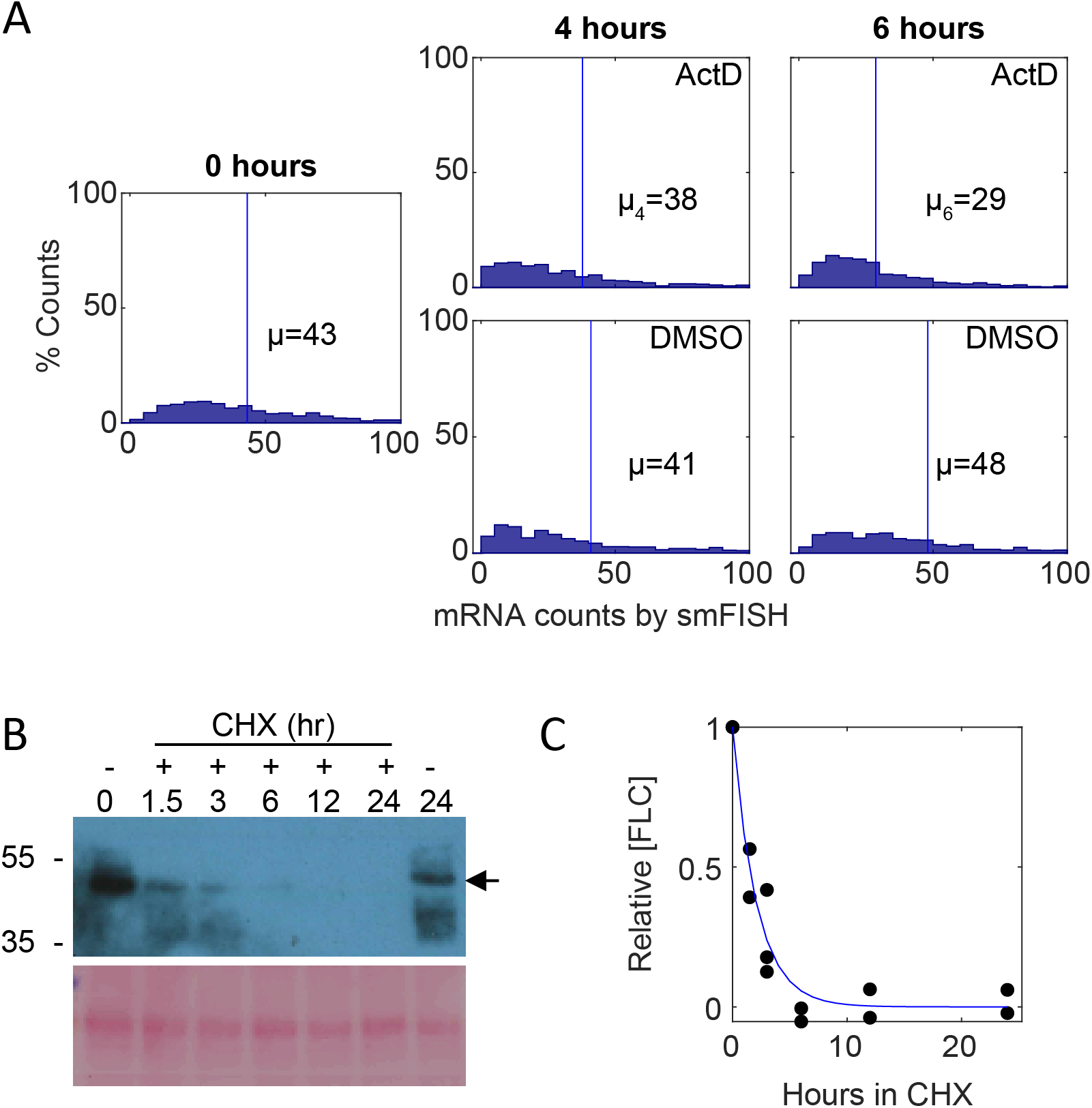
Effect of degradation rate and cell size variability on expected *FLC* levels. **(A)** Histogram of *FLC-Venus* mRNAs per cell measured by smFISH after 4 hours or 6 hours of ActD treatment (top) or mock DMSO treatment (bottom). Mean mRNA number per cell, µ, is indicated in each panel. Half-life t_1/2_ of 5 hours calculated as in (Ietswaart, Rosa et al. 2017) using the formula *t*_1/2_ = *ln*(2)/*d*, with *d* = ln (*μ*_4_/*μ*_6_)/2, where *d* is the degradation rate and *μ*_4_, *μ*_6_ are the mean RNA counts after 4 and 6 hours respectively. For 0 h, n=1554, for ActD 4 h, n=820, for ActD 6 h, n=784, for DMSO 4 h, n=759, for DMSO 6 h, n=776, in seedlings 7 days after sowing. **(B)** Analysis of FLC-Venus protein stability with cycloheximide (CHX) treatment. 7-day old seedlings of *fca-1* plants carrying FLC-Venus reporter were subjected to liquid MS solution supplemented with 100μM CHX for 0, 1.5, 3, 6, 12, or 24 hours. Blot shows representative replicate. Non treatment control was also included (right-most well). Arrow indicates FLC-Venus. Ponceau staining was used as processing controls. **(C)** Quantification of the FLC-Venus from blot shown in (B) and additional replicates. Combined data from n=3 biological replicates are shown with exponential decay profile fitted to first 3 timepoints (half-life=1.45hr).

**Figure 3 - Supplementary 2.**
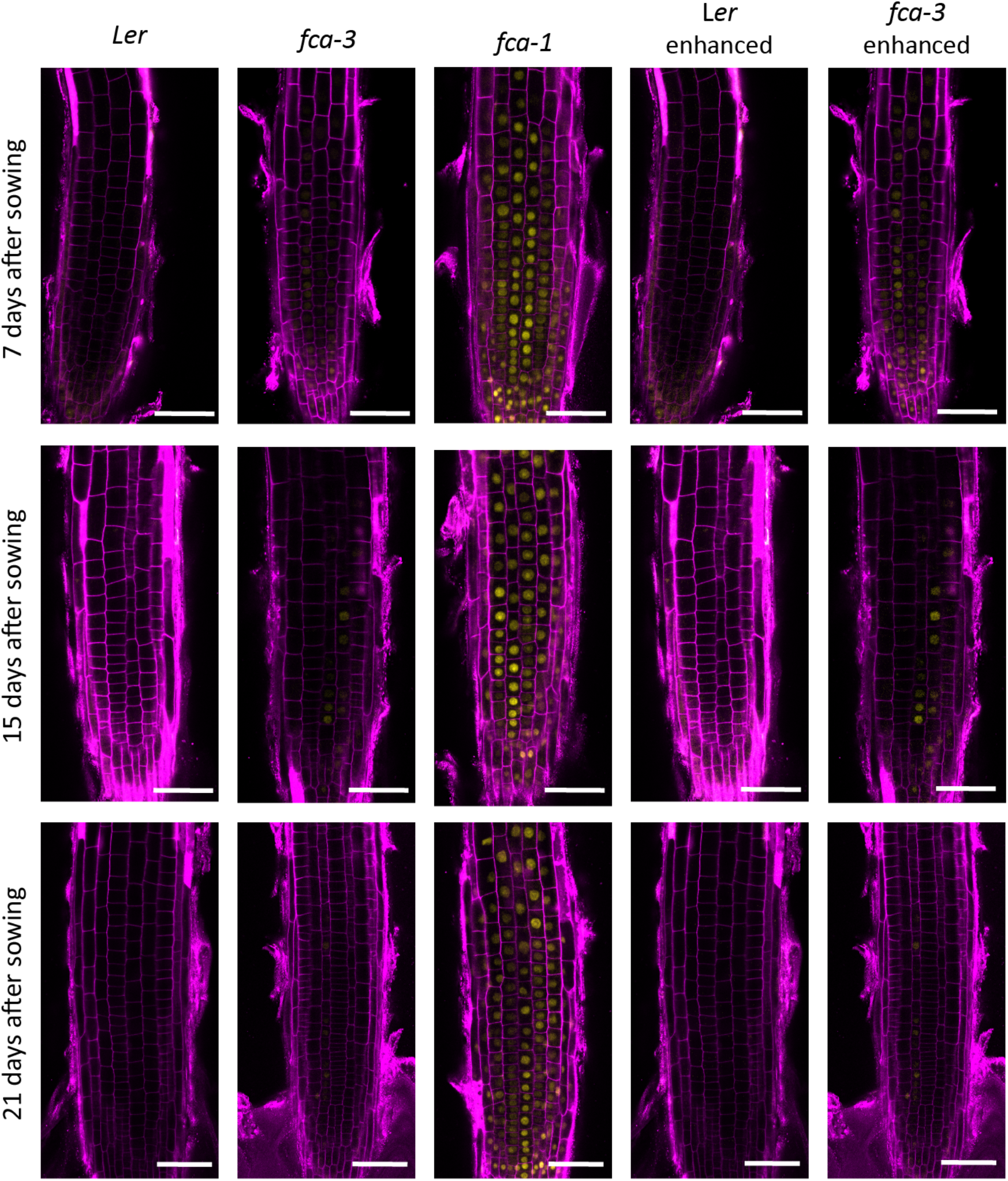
FLC-Venus timecourse replicates in *fca* alleles. Additional representative images of FLC-Venus signal in the three genotypes (Ler, *fca-3*, *fca-1*) at the three timepoints. Magenta shows the PI channel, Yellow is the FLC-Venus signal. Same settings were used for imaging and image presentation in left three columns. “Enhanced” images (two columns on right) are the same respective images, adjusted to enhance Venus signal to similar level as *fca-1* by changing brightness and contrast. Scale bar, 50 μm.

**Figure 3 - Supplementary 3.**
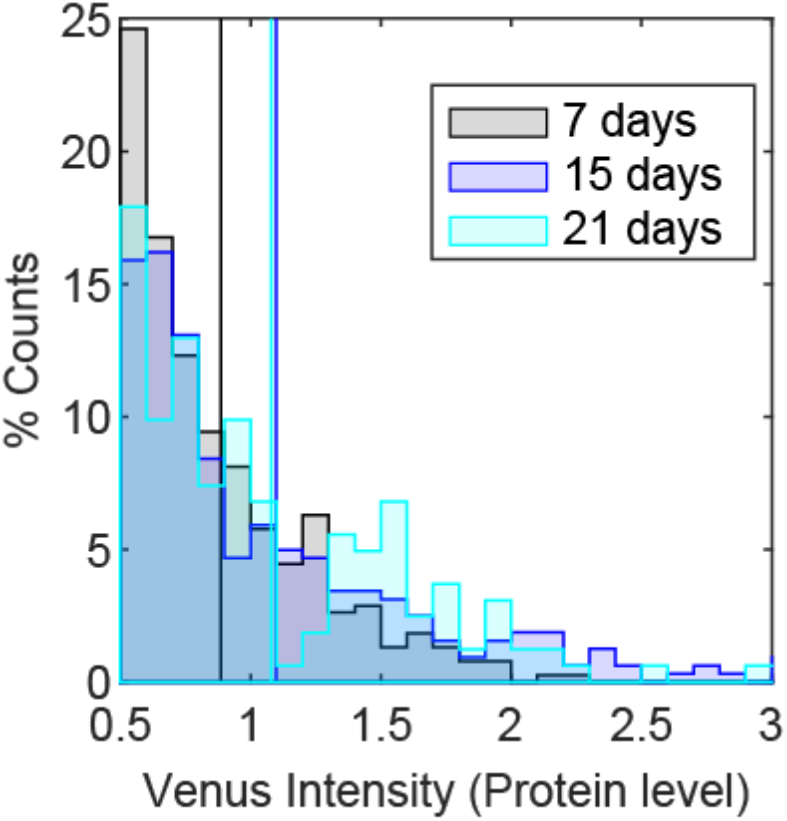
Experimental intensity of ON cells does not change over time in *fca-3*. Histogram of ON cells in *fca-3* at three timepoints normalised by the total number of ON cells in each. ON cells were defined as cells with FLC-Venus intensity above 0.5. Vertical lines indicate means of corresponding distribution. For 7 days n=382 cells from 24 roots, 15 days n=321 cells from 19 roots, 21 days n=162 cells from 12 roots.

**Figure 4 - Supplementary 1.**
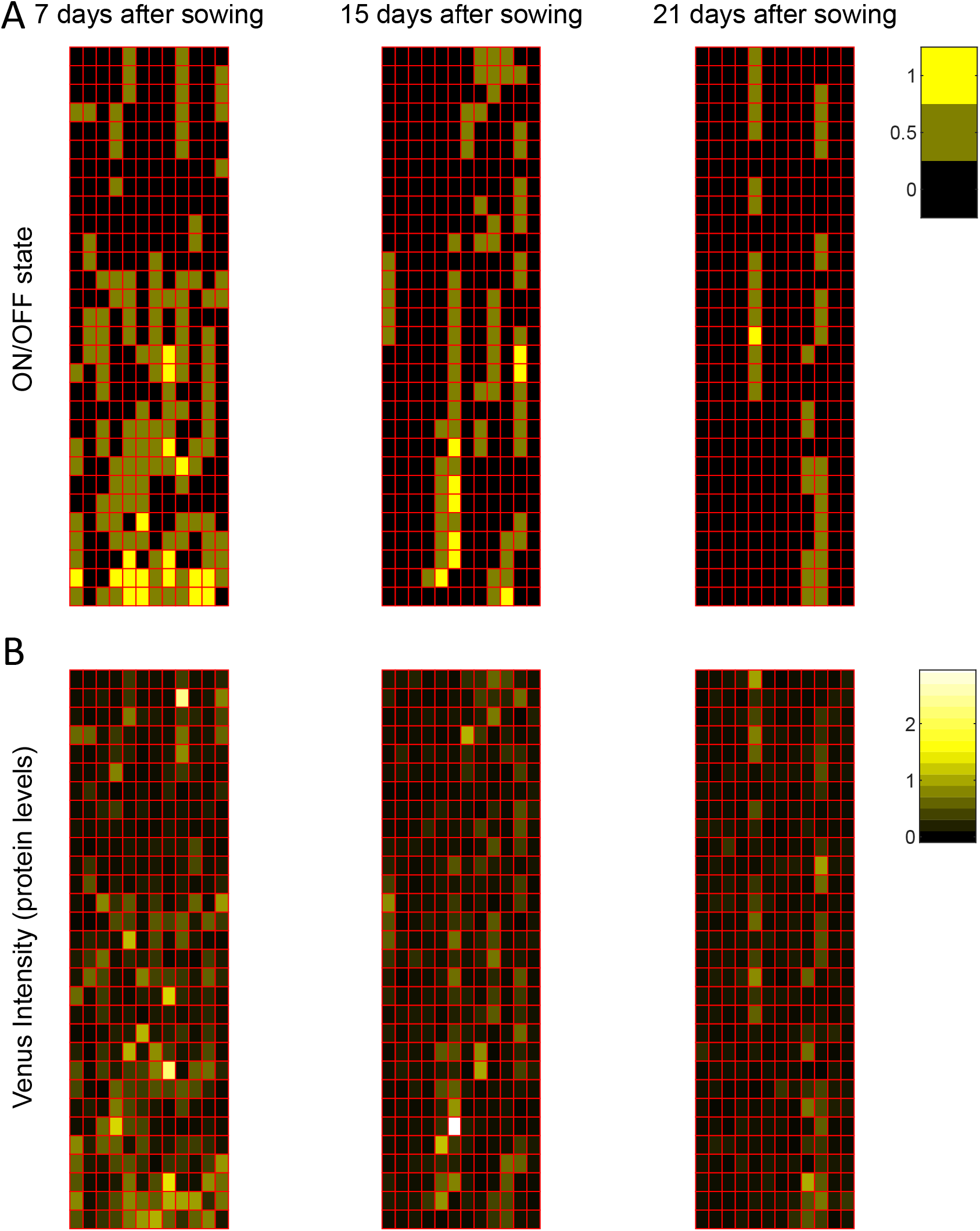
Modelling *FLC* regulation in clonal cell files in the root. Representative model output for *fca-3* root, showing 12 simulated cell files of length 30 cells each, at different timepoints, matching sampling times. **(A)** The state of the loci (both copies ON – yellow, one copy ON – grey, both copies OFF – black). **(B)** FLC-Venus intensity (proxy for protein concentration) per cell, for the same cells.

**Supplementary Table 1.**
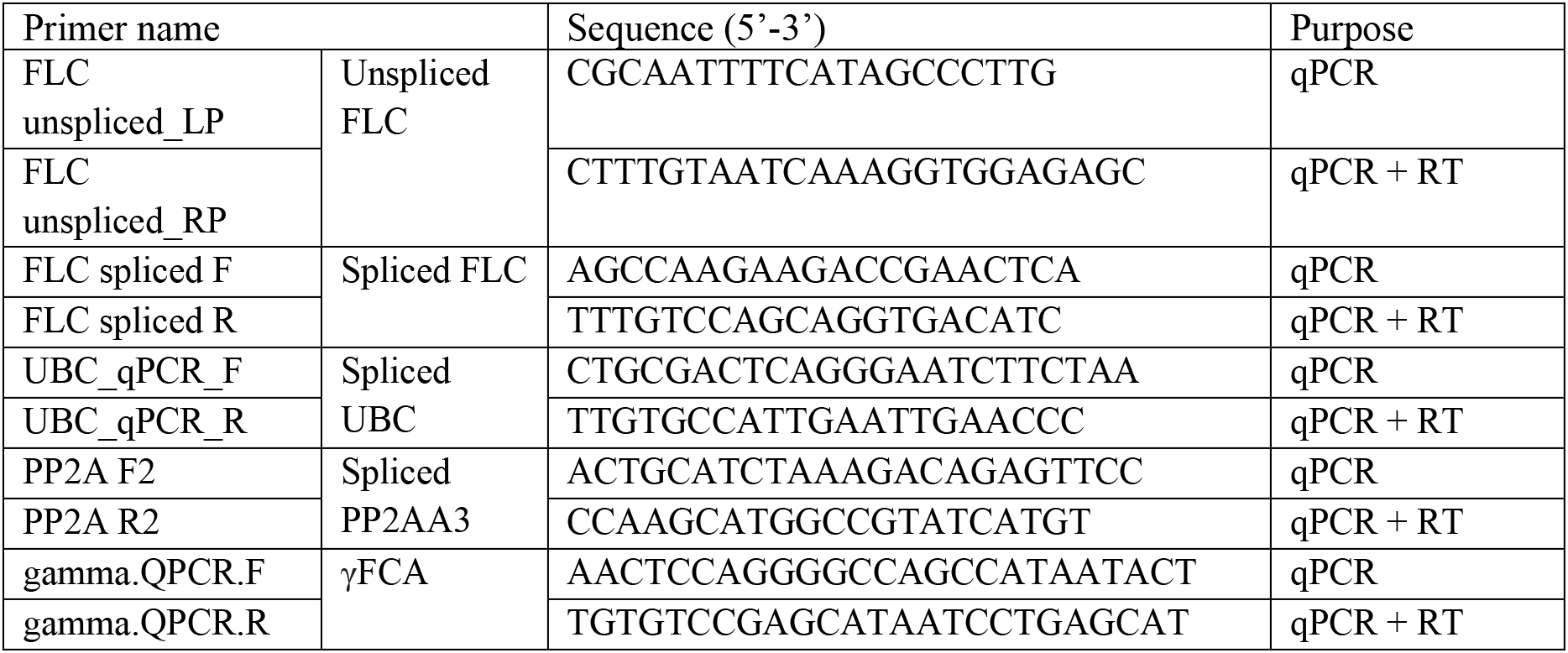
Primers used in this study.

**Supplementary Table 2:**
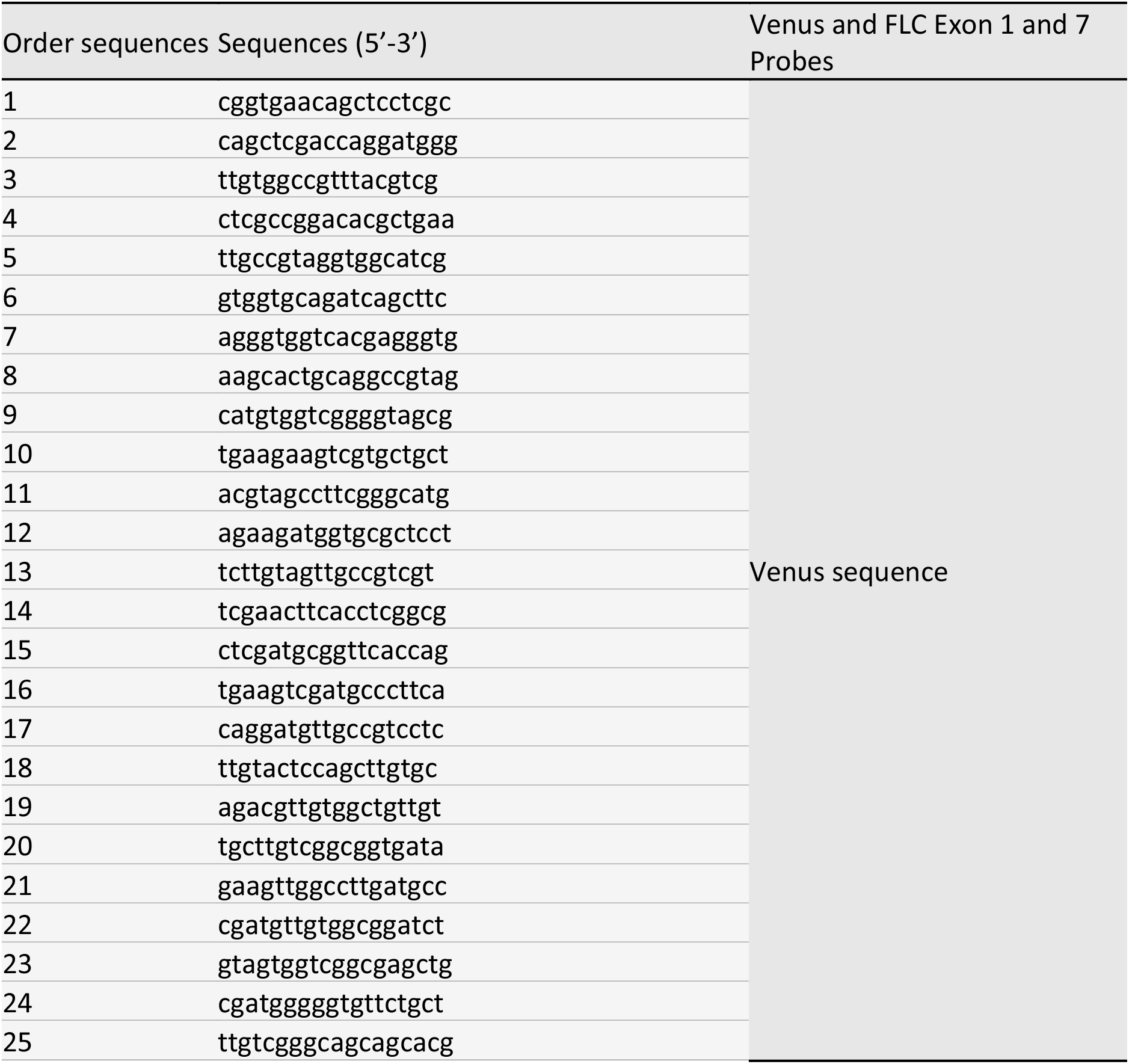
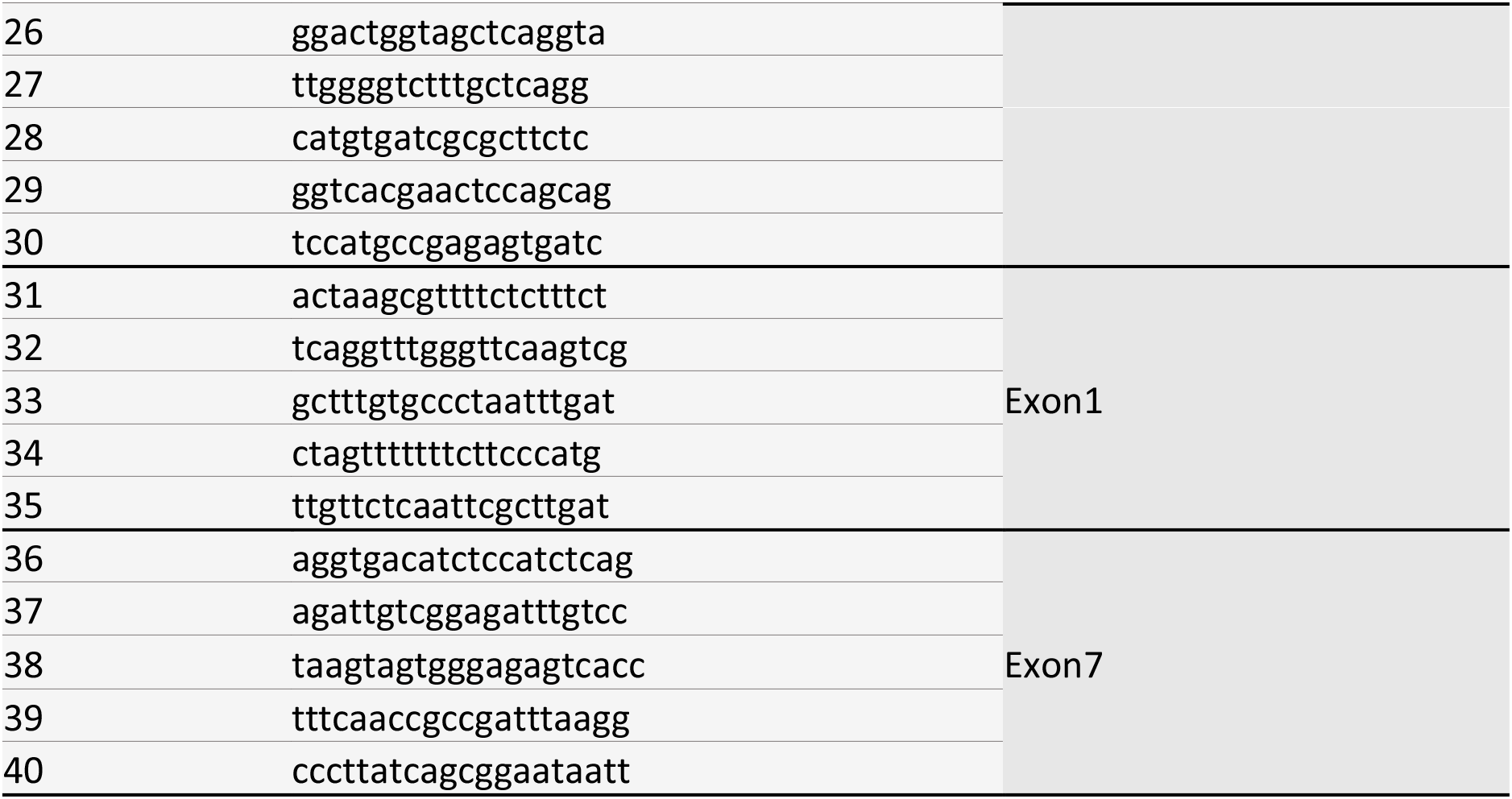
smFISH probe sequences used to detect FLC Venus transcripts. These probes were labelled with Quasar570.

**Supplementary Table 3:**
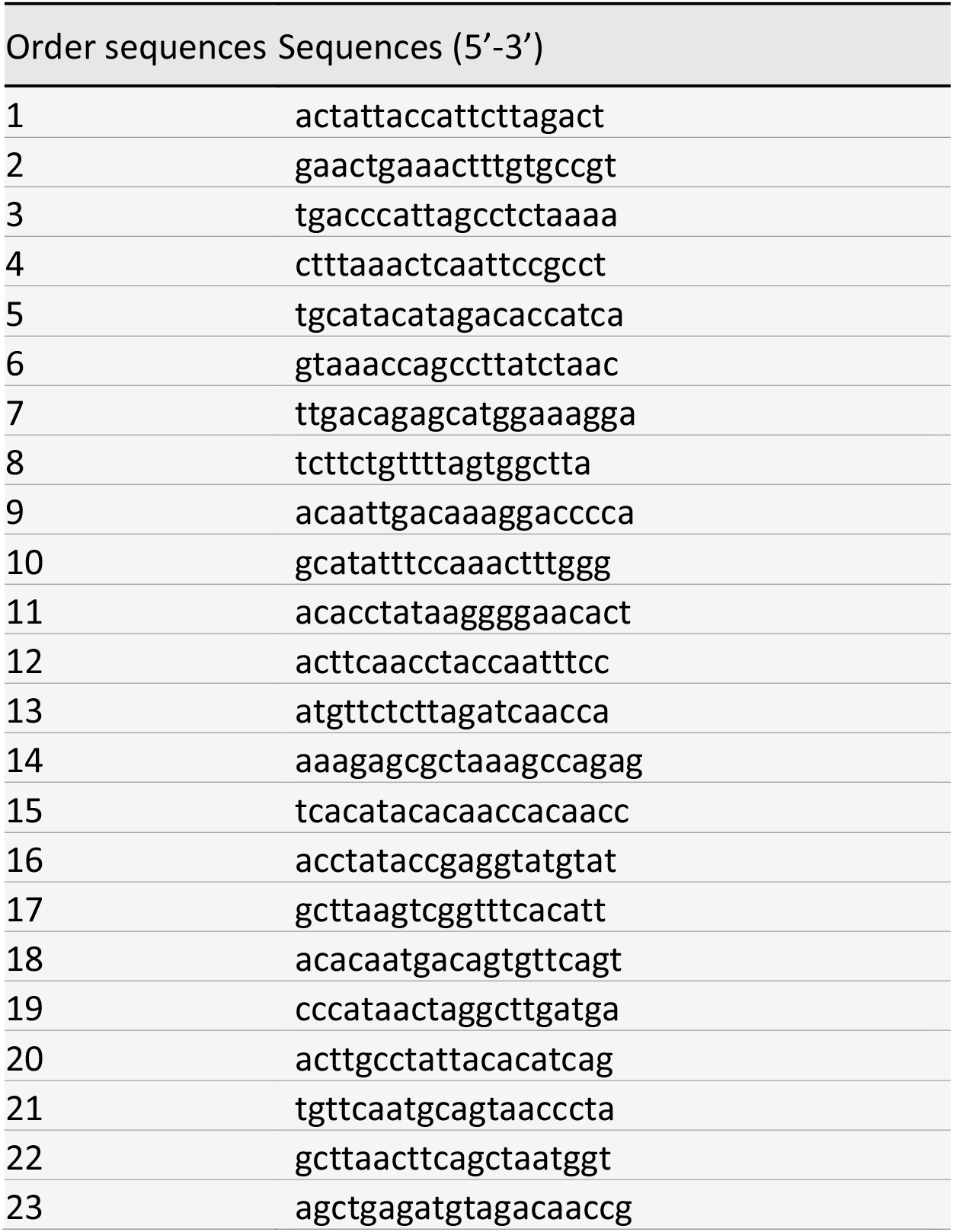
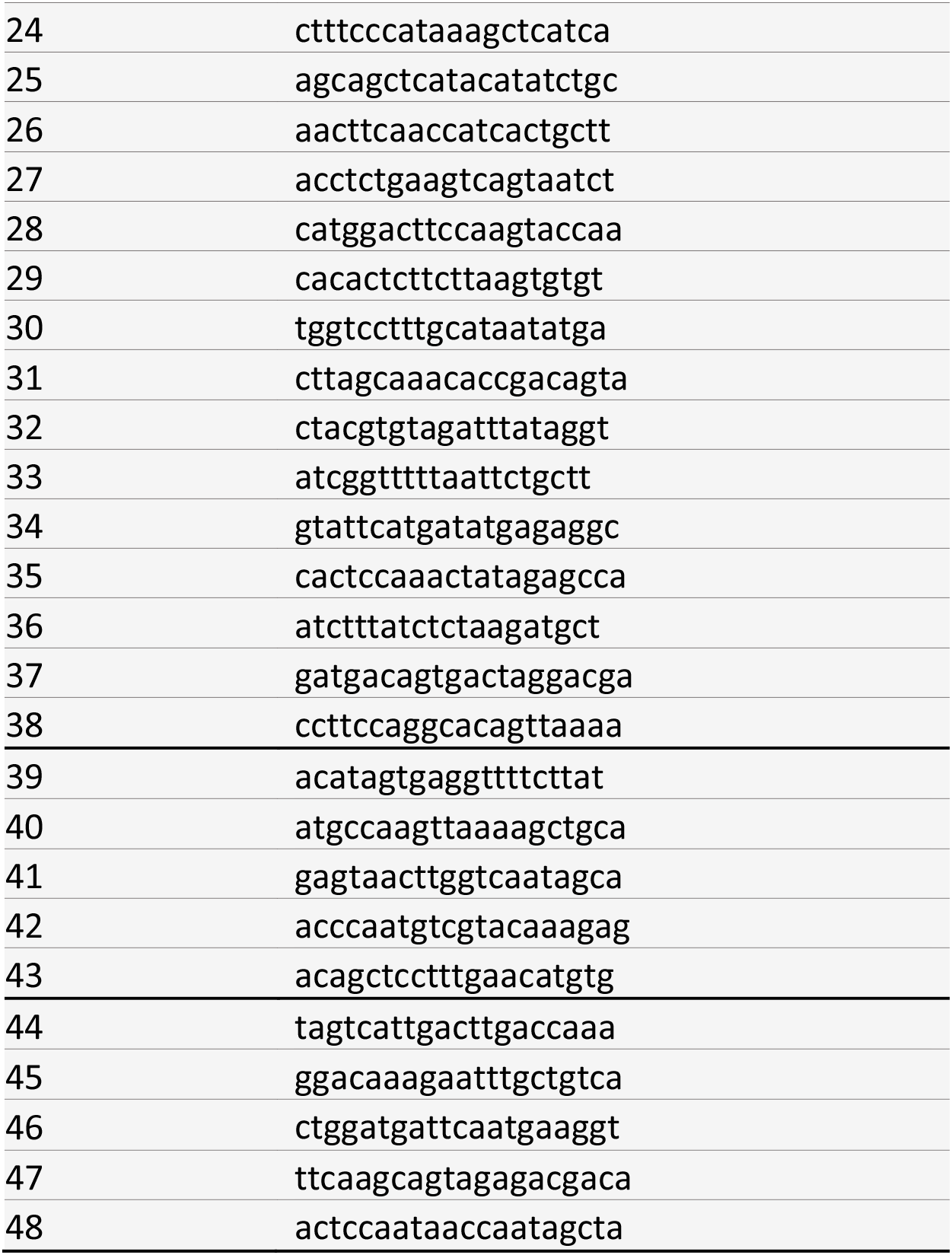
smFISH probe sequences used to detect unspliced PP2A transcripts. These probes were labelled with Quasar670.

**Supplementary Table 4.** Mathematical model parameters.

| Parameter name | Parameter value | Description |
| --- | --- | --- |
| p | 0.25 | Probability of a single *FLC* gene copy switching from ON to OFF during division. |
| $\mu_{ON}$ | -1.2 | $\mu$ of log normal distribution $e^X$, where $X \sim N(\mu, \sigma^2)$ for single ON *FLC* copy. |
| $\mu_{OFF}$ | -2.3 | $\mu$ of log normal distribution $e^X$, where $X \sim N(\mu, \sigma^2)$ for OFF *FLC* copy. |
| $\sigma_{ON}$ | 0.6 | $\sigma$ of log normal distribution $e^X$, where $X \sim N(\mu, \sigma^2)$ for ON *FLC* copy. |
| $\sigma_{OFF}$ | 0.4 | $\sigma$ of log normal distribution $e^X$, where $X \sim N(\mu, \sigma^2)$ for OFF *FLC* copy. |
| timestep | 1 hour | Simulation time step. |
| cell file length | 30 cells | Length of each simulated cell file, starting with initial cell (first cell after Quiescent Centre). |

## References

Ahmad, K., S. Henikoff and S. Ramachandran (2022). “Managing the Steady State Chromatin Landscape by Nucleosome Dynamics.” Annu Rev Biochem 91.

Angel, A., J. Song, C. Dean and M. Howard (2011). “A Polycomb-based switch underlying quantitative epigenetic memory.” Nature 476(7358): 105–108.

Baxter, C. L., S. Šviković, J. E. Sale, C. Dean and S. Costa (2021). “The intersection of DNA replication with antisense 3’ RNA processing in *Arabidopsis FLC* chromatin silencing.” Proceedings of the National Academy of Sciences 118(28): e2107483118.

Beare, R. and G. Lehmann (2006). “The watershed transform in ITK - discussion and new developments.” The Insight Journal 202: 1–24.

Beltran, M., C. M. Yates, L. Skalska, M. Dawson, F. P. Reis, K. Viiri, C. L. Fisher, C. R. Sibley, B. M. Foster, T. Bartke, J. Ule and R. G. Jenner (2016). “The interaction of PRC2 with RNA or chromatin is mutually antagonistic.” Genome Res 26(7): 896–907.

Berry, S. and C. Dean (2015). “Environmental perception and epigenetic memory: mechanistic insight through FLC.” Plant J.

Berry, S., C. Dean and M. Howard (2017). “Slow Chromatin Dynamics Allow Polycomb Target Genes to Filter Fluctuations in Transcription Factor Activity.” Cell Systems 4(4): 445–457.e448.

Berry, S., M. Hartley, T. S. G. Olsson, C. Dean and M. Howard (2015). “Local chromatin environment of a Polycomb target gene instructs its own epigenetic inheritance.” eLife 4: e07205.

Bintu, L., J. Yong, Y. E. Antebi, K. McCue, Y. Kazuki, N. Uno, M. Oshimura and M. B. Elowitz (2016). “Dynamics of epigenetic regulation at the single-cell level.” Science 351(6274): 720–724.

Coulon, A., M. L. Ferguson, V. de Turris, M. Palangat, C. C. Chow and D. R. Larson (2014). “Kinetic competition during the transcription cycle results in stochastic RNA processing.” eLife 3: e03939.

Gazzani, S., A. R. Gendall, C. Lister and C. Dean (2003). “Analysis of the molecular basis of flowering time variation in *Arabidopsis* accessions.” Plant Physiol. 132(2): 1107–1114.

Goodnight, D. and J. Rine (2020). “S-phase-independent silencing establishment in *Saccharomyces cerevisiae*.” eLife 9: e58910.

Holoch, D., M. Wassef, C. Lövkvist, D. Zielinski, S. Aflaki, B. Lombard, T. Héry, D. Loew, M. Howard and R. Margueron (2021). “A cis-acting mechanism mediates transcriptional memory at Polycomb target genes in mammals.” Nature Genetics 53(12): 1686–1697.

Ietswaart, R., S. Rosa, Z. Wu, C. Dean and M. Howard (2017). “Cell-size-dependent transcription of *FLC* and its antisense long non-coding RNA *COOLAIR* explain cell-to-cell expression variation.” Cell systems 4(6): 622–635. e629.

Johanson, U., J. West, C. Lister, S. Michaels, R. Amasino and C. Dean (2000). “Molecular analysis of FRIGIDA, a major determinant of natural variation in *Arabidopsis* flowering time.” <u>Science</u> 290(5490): 344–347.

Koornneef, M., C. J. Hanhart and J. H. v. d. Veen (1991). “A genetic and physiological analysis of late flowering mutants in *Arabidopsis thaliana*.” Molecular and General Genetics MGG 229: 57–66.

Kueh, H. Y., A. Champhekar, S. L. Nutt, M. B. Elowitz and E. V. Rothenberg (2013). “Positive Feedback Between PU.1 and the Cell Cycle Controls Myeloid Differentiation.” Science 341(6146): 670–673.

Lee, I. and R. M. Amasino (1995). “Effect of vernalization, photoperiod and light quality on the flowering phenotype of Arabidopsis plants containing the *FRIGIDA* gene.” Plant Physiol. 108: 157–162.

Li, Z., D. Jiang and Y. He (2018). “FRIGIDA establishes a local chromosomal environment for *FLOWERING LOCUS C* mRNA production.” Nature Plants 4(10): 836–846.

Lövkvist, C., P. Mikulski, S. Reeck, M. Hartley, C. Dean and M. Howard (2021). “Hybrid protein assembly-histone modification mechanism for PRC2-based epigenetic switching and memory.” eLife 10: e66454.

Michaels, S. D., Y. He, K. C. Scortecci and R. M. Amasino (2003). “Attenuation of *FLOWERING LOCUS C* activity as a mechanism for the evolution of a summer-annual flowering behavior in *Arabidopsis*.” Proc. Natl. Acad. Sci. U.S.A. 100(17): 10102–10107.

Munsky, B. and G. Neuert (2015). “From analog to digital models of gene regulation.” Phys Biol 12(4): 045004.

Python: Python Software Foundation, Python Language Reference, version 3.8.

Rahni, R. and K. D. Birnbaum (2019). “Week-long imaging of cell divisions in the Arabidopsis root meristem.” Plant Methods 15(1): 30.

Rosa, S., S. Duncan and C. Dean (2016). “Mutually exclusive sense-antisense transcription at FLC facilitates environmentally induced gene repression.” Nat Commun 7: 13031.

Saxton, D. S. and J. Rine (2022). “Distinct silencer states determine epigenetic states of heterochromatin.” bioRxiv: 2022.2002.2001.478725.

Schon, M., C. Baxter, C. Xu, B. Enugutti, M. D. Nodine and C. Dean (2021). “Antagonistic activities of cotranscriptional regulators within an early developmental window set *FLC* expression level.” Proceedings of the National Academy of Sciences 118(17): e2102753118.

Stewart-Ornstein, J., C. Nelson, J. DeRisi, J. S. Weissman and H. El-Samad (2013). “Msn2 coordinates a stoichiometric gene expression program.” Curr Biol 23(23): 2336–2345.

Wu, Z., X. Fang, D. Zhu and C. Dean (2019). “Autonomous Pathway: *FLOWERING LOCUS C* Repression through an Antisense-Mediated Chromatin-Silencing Mechanism.” Plant Physiology 182(1): 27–37.

Yang, H., M. Howard and C. Dean (2014). “Antagonistic roles for H3K36me3 and H3K27me3 in the cold-induced epigenetic switch at *Arabidopsis FLC*.” Curr Biol 24(15): 1793–1797.

Zhao, Y., R. L. Antoniou-Kourounioti, G. Calder, C. Dean and M. Howard (2020). “Temperature-dependent growth contributes to long-term cold sensing.” Nature 583(7818): 825–829.

